# Adaptation and Compensation in a Bacterial Gene Regulatory Network Evolving Under Antibiotic Selection

**DOI:** 10.1101/2021.08.31.458319

**Authors:** Vishwa Patel, Nishad Matange

## Abstract

Gene regulatory networks allow organisms to generate coordinated responses to environmental challenges. In bacteria, regulatory networks are re-wired and re-purposed during evolution, though the relationship between selection pressures and evolutionary change is poorly understood. In this study, we discover that early evolutionary response of *Escherichia coli* to the antibiotic trimethoprim involves de-repression of PhoPQ signalling, a Mg^2+^-sensitive two-component system, by inactivation of the MgrB feedback-regulatory protein. We report that de-repression of PhoPQ confers trimethoprim-tolerance to *E. coli* by hitherto unrecognized transcriptional up-regulation of dihydrofolate reductase (DHFR), target of trimethoprim. As a result, mutations in *mgrB* precede and facilitate the evolution of drug resistance. Using laboratory evolution, genome sequencing and mutation re-construction, we show that populations of *E. coli* challenged with trimethoprim are faced with the evolutionary ‘choice’ of transitioning from tolerant to resistant by mutations in DHFR, or compensating for the fitness costs of PhoPQ de-repression by inactivating the RpoS sigma factor, itself a PhoPQ-target. Outcomes at this evolutionary branch-point are determined by strength of antibiotic selection, such that high pressures favour resistance, while low pressures favour cost-compensation. Our results relate evolutionary changes in bacterial gene regulatory networks to strength of selection and provide mechanistic evidence to substantiate this link.

## Introduction

The relationship between genotype and phenotype is complex and much of modern genetics aims at understanding how it is established and maintained. In cellular organisms, this relationship is dictated in large part by regulation of gene expression in response to internal and external environment [1–3]. By regulating gene expression, organisms select which traits are expressed, enabling a wide palette of phenotypes to be generated from the same genotype. Not surprisingly, the functioning and evolution of mechanisms that regulate gene expression are central questions at the cusp of genetics and evolutionary biology with implications for diverse phenomena such as development [4], differentiation [4] and disease [5].

In bacteria, like in more complex organisms, gene regulatory proteins, such as transcription factors and signal transducers are organized into networks [6]. The predominant sensory modules in bacteria are two-component signalling systems, which are made up of membrane-bound receptors and cognate cytosolic response regulators [7]. Upon activation of the receptor by an appropriate ligand, a phospho-relay between the two components is initiated, which changes the phosphorylation state of the response regulator. Response regulators are often transcription factors themselves, which bind to DNA upon phosphorylation and produce changes in gene expression [7]. Individual two-component pathways are integrated into networks by ‘connector’ proteins, and also by cross-activation [8]. Further, two-component systems are integrated with other signalling molecules in bacteria such as regulatory RNAs [9] and Sigma factors [10].

Gene regulatory networks orchestrate a number of physiological functions in bacteria such as sporulation [11], virulence [12], dormancy [13] and tolerance to stresses like acid [14], heavy metal [15] and osmolarity [16]. More recently, the role of bacterial gene regulatory networks in antimicrobial resistance is beginning to emerge [17]. VanRS (vancomycin resistance, *Staphylococcus aureus*) [18], BaeRS (ciprofloxacin resistance, *Salmonella typhimurium*) [19], PhoPQ [20, 21] and PmrAB [22] (antimicrobial peptide resistance, *Salmonella* and *Escherichia coli*), MtrAB (multidrug resistance, *Mycobacterium tuberculosis*) [23] and CroRS (β-lactam resistance, *Enterococcus faecalis*) [24] are some of the two-component systems that mediate drug resistance in bacteria. Two-component systems modulate antibiotic resistance by regulating the expression of down-stream effectors such as efflux pumps, porins or enzymes that modify the cell envelope [17]. Not surprisingly, two component sensor kinases have also been proposed as potential targets for the design of novel anti-microbials [25].

The repertoire of two-component systems that bacteria possess depends on environment and life-history. For instance, *Myxococcus xanthus* that faces rapidly-changing, diverse environments codes for more than 250 different two-component systems [26]. Further, rewiring of two-component systems during evolution is also seen among bacteria. For example, the regulons of Mg^2+^-sensitive PhoPQ systems from closely related members of *Enterobacteriaceae* show several species-specific features. Most notably, *Salmonella* and *Yersinia* have only a marginal overlap in PhoP-regulons, even though they share a common ancestor [27]. Similarly, co-option of existing regulatory modules for the expression of genes coded by horizontally-acquired pathogenicity islands in *Salmonella* has also been described [28]. Thus, two-component systems and their associated transcriptional networks are likely to be ready substrates for evolution to suit the unique demands of different growth environments.

Current understanding of the evolution of two-component signalling systems and their associated gene regulatory networks comes from two main approaches. First of these relies on comparison of orthologous genes from related bacterial species [27, 29]. The second uses genetically-engineered bacterial strains to understand general principles governing the specificity of receptor-regulator pairs and the potential for cross-activation in two-component systems [30]. While invaluable insight has emerged from both approaches, the direct link between evolutionary changes in gene regulatory networks and selection pressures remains elusive. In this study, we use laboratory evolution to identify a specific environmental factor, i.e. the antibiotic trimethoprim, as a selective pressure that leads to de-repression of the PhoPQ signalling pathway in *E. coli.* This adaptation is mediated by loss-of-function mutations at the *mgrB* locus, which codes for a feedback attenuator of PhoPQ signalling. Intrinsic resistance to trimethoprim in *E. coli* is thought to be acquired primarily by mutations in the drug-target dihydrofolate reductase (DHFR), coded by the *folA* gene [31, 32]. We find that mutations in DHFR are typically preceded by mutations at the *mgrB* locus and together, they have a synergistic effect of the level of trimethoprim resistance. We use this system to ask how the costs of adaptation by perturbation of gene regulatory networks are compensated over long-term evolution and identify the key role that strength of selection plays in determining the adaptive trajectory that bacteria take upon antibiotic challenge.

## Results

### Mutation at the *mgrB* locus rather than *folA* is an early evolutionary response of *E. coli* challenged with trimethoprim

We had earlier evolved trimethoprim-resistance in *E. coli* K-12 MG1655 by serially passaging bacteria in drug-supplemented medium for ~25 generations [33]. From these experiment 10 independently evolved trimethoprim-resistant isolates (designated TMPR1-10), with drug IC50 ranging from 6- to 300-fold over wild type were obtained (Figure 1A). Unexpectedly, only 3 of the 10 trimethoprim-resistant isolates (TMPR 1, 3 and 5) had mutations at the *folA* locus (promoter and coding region), which suggested that early adaptation to trimethoprim did not necessarily involve mutations in the drug target itself (Figure 1A). We, therefore, sequenced the genomes of 5 of the 10 isolates (TMPR1-5) to identify other genes that may contribute to trimethoprim resistance. Only one locus, coding for the *mgrB* feedback regulator protein, consistently harboured mutations in all 5 isolates (Table 1). Four of the 5 isolates (TMPR1-4) harboured IS-element insertions in the promoter of *mgrB*. These insertions are expected to reduce expression of *mgrB* by preventing transcriptional activation by PhoP (Figure 1A, Table 1). The fifth isolate (TMPR5) harboured a single nucleotide deletion in the coding region of *mgrB* that results in stop-codon readthrough and altered sequence of the C-terminal domain of the protein (Figure 1A, S5, Table 1). The C-terminus of MgrB is necessary for its inhibitory activity against PhoQ [34]. We therefore argued that lower MgrB expression or activity may be advantageous for *E. coli* in the presence of trimethoprim. We tested this hypothesis using an independently generated *ΔmgrB* strain and found that loss of MgrB enhanced trimethoprim IC50 by ~3-fold (Figure 1B, C). Thus, mutation-driven loss of *mgrB*, rather than mutations in the *folA* gene, was the predominant early adaptive change in *E. coli* challenged with trimethoprim.

**Figure 1.**
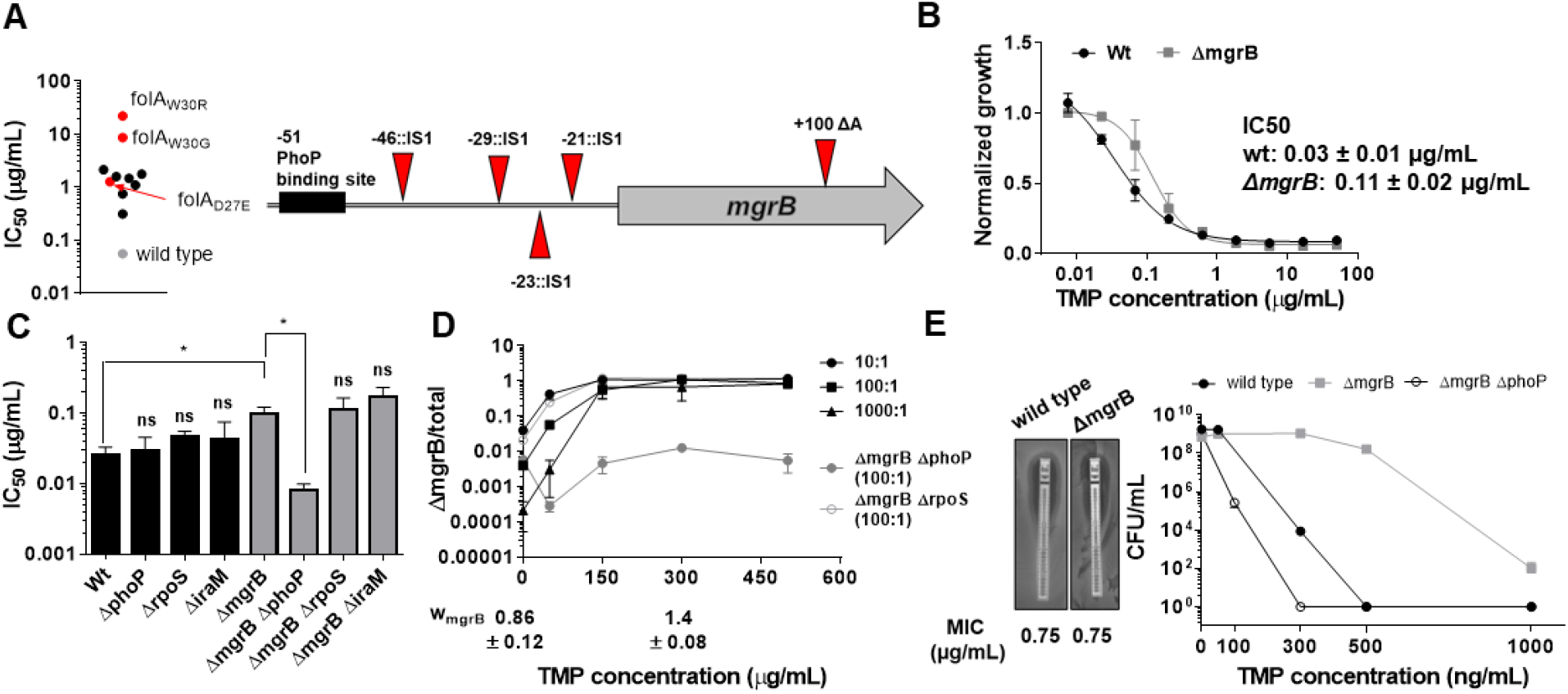
Loss of *mgrB* confers trimethoprim tolerance to *E. coli*. **A**. Left panel. Trimethoprim IC50 values of resistant isolates derived from *E. coli* K-12 MG1655 (wild type, gray) by laboratory evolution. Mean IC50 values from two independent measurements are plotted. Isolates without mutations in the *folA* locus are represented by black circles. Isolates with mutations in *folA* are represented as red circles, and the identified mutation is indicated. Right panel. Diagrammatic representation of the *mgrB* gene and its promoter showing mutations identified in trimethoprim resistant isolates by genome sequencing. The PhoP-binding site in the *mgrB* promoter is shown as a black box. Location of each mutation is calculated as base pairs from the translation start site. **B**. Growth of wild type (black) and *ΔmgrB* (gray) *E. coli* in varying concentrations of trimethoprim, normalized to growth in drug-free medium. Each data point represents mean ± S.D. from 3 independent experiments. IC50 values represent mean ± SEM obtained after curve fitting. **C**. IC50 values of trimethoprim for wild type or mutant *E. coli.* Each bar represents mean ± SEM from 3 independent measurements. Statistical significance was tested using a Student’s t-test. A p-value of <0.05 was considered significant (*). **D**. Competition between *E. coli* wild type and *ΔmgrB* in increasing concentrations of trimethoprim starting at the indicated mixing ratios (black). The fraction of *ΔmgrB* bacteria in each mixed culture after ~9 generations of competition are plotted (mean ± SD from 3 independent experiments). Values of relative fitness for *E. coli ΔmgrB* (w_mgrB_) at 0 and 300 ng/mL trimethoprim are given below the graph. Results of competition between wild type and *ΔmgrBΔrpoS* (open circles) or *ΔmgrBΔphoP* (solid circles) strains at an initial mixing ratio of 100:1 are shown in gray. **E**. Left panel. MIC values of trimethoprim for wild type and *E. coli ΔmgrB* calculated from E-tests. Right panel. Colony formation of wild type and *ΔmgrB* on solid media supplemented with indicated concentrations of trimethoprim. Each point represents mean ± S.D. from 3 independent experiments.

**Table 1.** Summary of mutations in trimethoprim resistant isolates TMPR1-5 identified by genome sequencing.

| Locus | Description | TMPR1 | TMPR2 | TMPR3 | TMPR4 | TMPR5 |
| --- | --- | --- | --- | --- | --- | --- |
| Single nucleotide changes |  |  |  |  |  |  |
| folA | dihydrofolate reductase | Asp27Glu | - | Trp30Gly | - | Trp30Arg |
| ycaQ | inter-strand DNA crosslink repair glycosylase | - | - | Trp71* | - | - |
| yagF | D-xylonate dehydratase | - | - | - | - | Ser557Pro |
| yagH | putative xylosidase/arabinosidase | - | - | - | - | Val397Met |
| mgrB | PhoQ kinase inhibitor | - | - | - | - | Deletion +100::ΔA |
| hexR | DNA-binding transcriptional repressor | - | - | - | - | Ala143Val |
| deoA | thymidine phosphorylase | Insertion +359::A | - | Insertion +375::A |  |  |
| dnaA | chromosomal replication initiator protein | - | - | - | Asp280Gly | - |
| Insertion elements and large rearrangements |  |  |  |  |  |  |
| mcrC | 5-methylcytosine-specific restriction enzyme subunit | - | - | +969::IS1A | - | - |
| ybcM | putative DNA-binding transcriptional regulator | +493::IS1A | - | - | - | - |
| mgrB | PhoQ kinase inhibitor | -23::IS1A | -29::IS1A | -46::IS1A | -21::IS1A | - |
| Partial genome duplication | - | - | - | - | 2X (~1 to 7,00,000) | - |

### Loss of *mgrB* confers trimethoprim tolerance by PhoP-dependent up-regulation of DHFR expression

To better understand how loss of *mgrB* reduced the effectiveness of trimethoprim against *E. coli*, we characterized the *mgrB* knock-out strain. *E. coli ΔmgrB* had lower relative fitness than wild type in antibiotic-free media (Figure 1D). However, sub-MIC (minimum inhibitory concentration) trimethoprim in growth media enhanced relative fitness of the mutant (Figure 1D). Concomitantly, *E. coli ΔmgrB* was selected over wild type in the presence of trimethoprim, even when initial mixing ratios were biased in favour of the latter (Figure 1D). There was no detectable difference in trimethoprim MIC between wild type and mutant (Figure 1B, E). However, *mgrB*-deficient *E. coli* formed colonies even at concentrations close to MIC, albeit at significantly reduced efficiency (Figure 1E). Further, relatively fewer cells of *E. coli ΔmgrB* were required to colonize growth media with inhibitory concentrations of trimethoprim (Figure S1). These experiments established that loss of *mgrB* allowed better survival of *E. coli* over a range of trimethoprim concentrations. However, since no change in drug MIC was detected, we concluded that *mgrB*-deficiency conferred trimethoprim tolerance, rather than resistance, to *E. coli*.

The primary role of the MgrB protein in *E. coli* is to attenuate PhoPQ signalling through negative feedback [35, 36]. Thus, PhoPQ signalling is de-repressed in an *mgrB* knockout strain [35, 36]. In line with this role, deletion of *phoP* re-sensitized *E. coli ΔmgrB* to trimethoprim, while in a wild type background had no detectable effect (Figure 1C, D, E, S1). PhoP directly activates transcription of close to 50 genes in *E. coli* with diverse functions [37]. Among these is *iraM*, which protects the stress-responsive sigma factor RpoS from degradation [38]. Arguing that trimethoprim tolerance in the *mgrB-*knockout could be a result of enhanced general stress-response pathway, we generated knockout strains of *iraM* and *rpoS* in a *ΔmgrB* background. However, neither deletion altered trimethoprim susceptibility of *ΔmgrB*, ruling-out this possibility (Figure 1C, D, E, S1).

To identify the target of PhoP that led to trimethoprim-tolerance, we turned to the transcriptomics data published by Xu et al. (2019) [38] that compared gene expression profiles of wild type and *mgrB-*deficient *E. coli.* Among over 500 genes up-regulated in *mgrB*-deficient *E. coli*, 97 are known targets of RpoS and hence could be excluded (Figure S2). Curiously, *folA* transcript levels were reported to be enhanced by ~3-fold in an *mgrB*-deficient strain [38]. The *folA* gene codes for DHFR, target of trimethoprim, and its overexpression results in trimethoprim resistance [39]. We found that DHFR protein levels were indeed elevated in an *mgrB* knockout strain (Figure 2A). Further, DHFR levels could be lowered in the *mgrB*-knockout by deletion of*phoP* or *phoQ*, but not *rpoS* or *iraM* (Figure 2B). The increase in DHFR levels was due to greater activity of the *folA* promoter in *E. coli ΔmgrB* compared to wild type (Figure 2B, C), and this too was *phoP*-dependent but *rpoS*-independent (Figure 2C). Importantly, the *folA* promoter retained trimethoprim-stimulation in the *mgrB*-knockout (Figure 2C). The *folA* promoter is not a known target for PhoP, and does not harbour sequences similar to the PhoP binding site. Consequently, we were unable to detect direct binding of PhoP to the *folA* promoter *in vitro* by gel shift assays (Figure S3). Thus, the enhancement of *folA* expression by PhoP was most probably an indirect effect.

**Figure 2.**
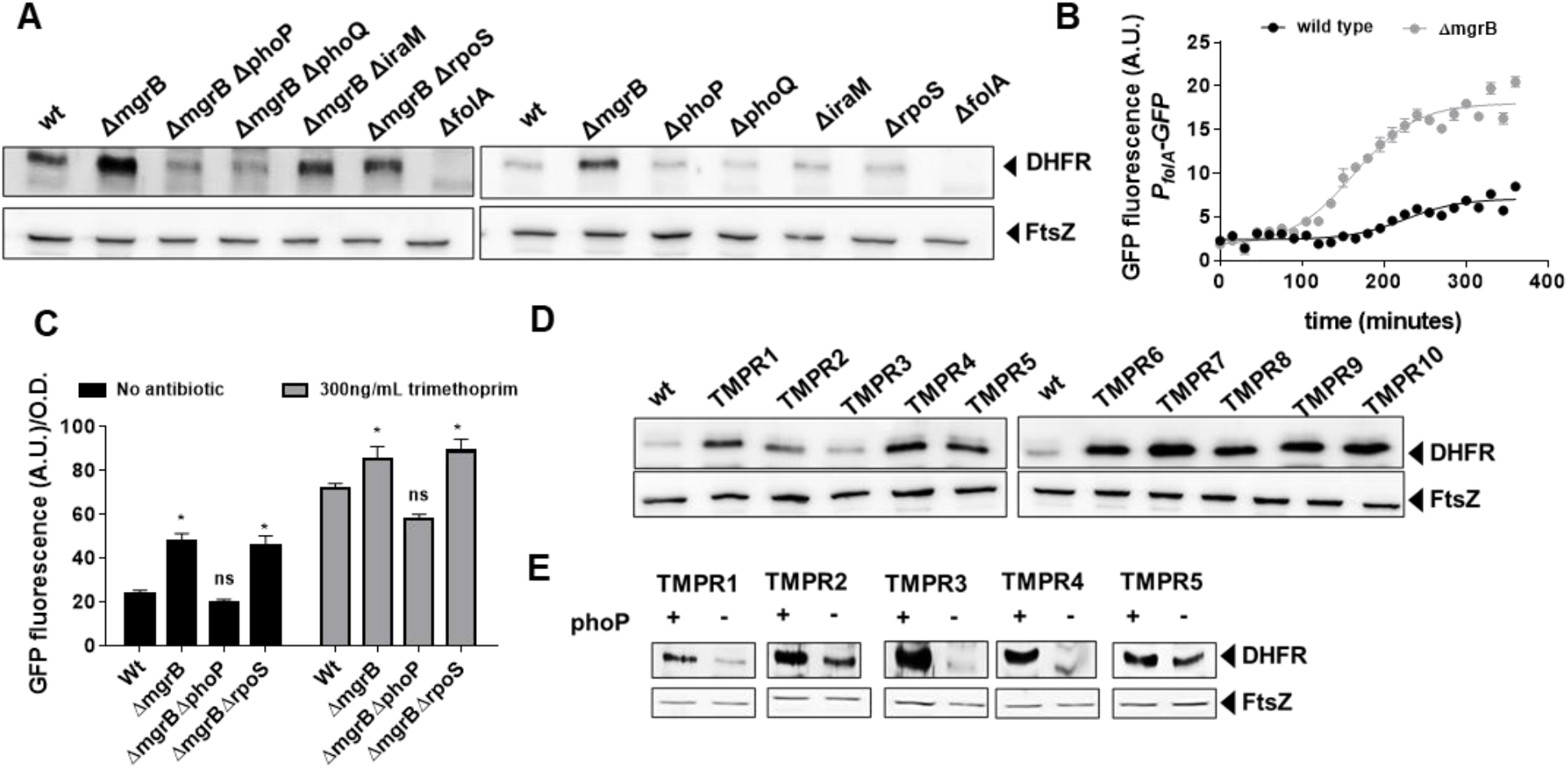
Enhanced expression of DHFR in *mgrB-*deficient *E. coli*. **A.** Lysates (5 μg total protein) of wild type or mutant *E. coli* subjected to immunoblotting using anti-DHFR polyclonal antibody. FtsZ was used as loading control. Data shown are representative of 3 independent replicates. **B**. Activity of the *folA* promoter (P_folA_.) in *E. coli* wild type or *ΔmgrB* monitored over growth using a GFP reporter gene (arbitrary units, A.U.). Each point represents mean ± S.D. from 3 replicates. **C**. Peak fluorescence normalized to optical density for indicated strains harbouring the P_fol_-GFP reporter plasmid. Mean ± S.D. from 3 independent experiments is plotted. Statistical significance was tested using the Student’s t-test and a p-value of < 0.05 was considered significant (*). Promoter activity was measure in drug-free medium or in the presence of 300 ng/mL trimethoprim. **D**. Lysates (5 μg total protein) of *E. coli* wild type or trimethoprim resistant isolates (TMPR1-10) subjected to immunoblotting using anti-DHFR polyclonal antibody. FtsZ was used as loading control. Data shown are representative of 3 independent replicates. **E**. Lysates (5 μg total protein) of trimethoprim-resistant isolates (TMPR1-5) and their *ΔphoP* derivatives subjected to immunoblotting using anti-DHFR polyclonal antibody. FtsZ was used as loading control. Data shown are representative of 3 independent replicates.

In line with higher expression of DHFR in *mgrB*-deficient *E. coli*, 9 of the 10 trimethoprim-resistant isolates (TMPR1-2,4-10) showed elevated DHFR protein levels (Figure 2D). The only isolate that did not (TMPR3), harboured the W30G mutation in *folA*, which we have shown earlier to result in lower steady state levels of DHFR. Further, deletion of *phoP* resulted in lower expression of DHFR in TMPR1-5 (Figure 2E). These data demonstrated that loss of *mgrB* conferred trimethoprim tolerance in *E. coli* by enhancing DHFR protein levels, through PhoP-dependent, RpoS-independent transcriptional up-regulation of the *folA* promoter.

### De-repression of PhoPQ facilitates resistance evolution by altering the fitness landscape of *folA* mutations

In addition to *mgrB* mutations, isolates TMPR1, 3 and 5 harboured mutations in *folA* (Table 1). Likewise, TMPR4 harboured a large genomic duplication encompassing the *folA* gene along with other genes that may confer trimethoprim resistance (Table 1). To assess the contribution of loss of *mgrB* to the phenotypes of these isolates, we tested trimethoprim MIC and IC50 of *ΔphoP* derivative of TMPR1-5. Remarkably, loss of *phoP* greatly reduced the MIC of trimethoprim for all 5 isolates (Figure 3A). TMPR2 and 4 that did not harbour mutations in *folA* were completely re-sensitized to trimethoprim by *phoP* deletion (Figure 3A, B). TMPR1, 3 and 5, which harboured mutations in *folA* retained some trimethoprim-resistance after deletion of *phoP*, but with close to 10-fold reduction in drug IC50 (Figure 3B). Thus, even though loss of *mgrB* itself did not appreciably alter drug MIC, it significantly potentiated the phenotypes of *folA* mutations in trimethoprim-resistant *E. coli*.

**Figure 3.**
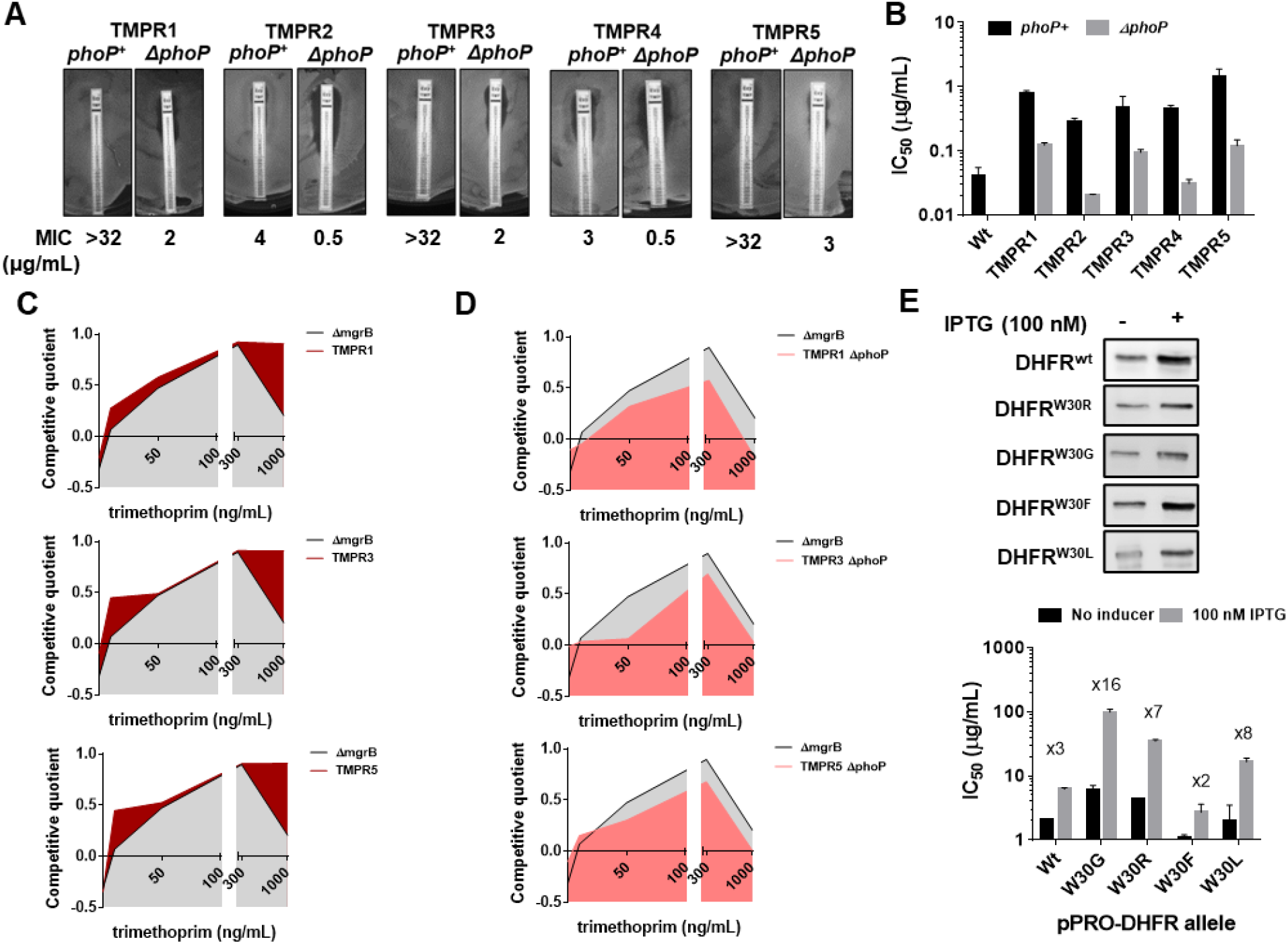
Synergy between *mgrB* and *folA* mutations alters the fitness landscape of trimethoprim resistant *E. coli*. **A.** MIC values of trimethoprim for resistant isolates (TMPR1-5) and their *ΔphoP* derivatives calculated from E-tests. **B**. IC50 values of trimethoprim from resistant isolates (TMPR1-5) and their *ΔphoP* derivatives. Mean ± S.E.M from 3 independent experiments are plotted. **C, D**. Fitness landscape of trimethoprim resistant isolates TMPR1, 3 and 5 (C) or their *ΔphoP* derivatives (D) compared with trimethoprim tolerant *E. coli ΔmgrB*. Competitive quotients were calculated by competition with *E. coli-*GFP (wild type *E. coli* expressing GFP) in varying concentrations of trimethoprim. Higher values indicate higher fitness relative to wild type. Representative data from three replicates are shown. **E**. Effect of overexpression of DHFR on IC50 of trimethoprim monitored by expression of wild type or mutant DHFR from an IPTG-inducible promoter. Upper panel. Immunoblots showing roughly similar over-expression of all DHFR alleles tested. Lower panel. IC50 values of trimethoprim for various mutants with and without inducer are plotted (mean ± S.D. from 3 replicates). The fold increase in IC50 in the presence of inducer is indicated for each mutant.

The above result prompted us to ask whether mutation at *mgrB* influenced the selection dynamics of trimethoprim resistance in *E. coli*. The genetic constitution of antibiotic resistant bacteria is known to be influenced by drug concentration [40–42]. Hence, we mapped the fitness landscape of drug-tolerant (*E. coli ΔmgrB*) and drug-resistant (TMPR1, 3 and 5) strains across different concentrations of trimethoprim (Figure 3C). Each of the above strains was allowed to compete against a GFP-expressing derivative of *E. coli* wild type (*E. coli-*GFP). Since GFP fluorescence of the mixed culture would indicate the fractional abundance of *E. coli-GFP* in the population, we used it as a read-out of relative fitness of the test strains (designated as competitive quotient; a value of 0 indicated no change in fitness relative to wild type, >0 signified higher relative fitness, <0 signified reduced relative fitness). Over the entire range of trimethoprim concentrations used by us, TMPR1, 3 and 5 performed better than *ΔmgrB* alone. This difference was most pronounced at concentrations that approached the MIC value for the wild type. Next, we asked how *ΔphoP* derivatives of TMPR1, 3 and 5 would perform in a similar assay (Figure 3D). Interestingly, *ΔmgrB* was fitter than *ΔphoP* derivatives of TMPR1, 3 and 5 across all concentrations of trimethoprim tested by us. These results explained why *mgrB* mutations were the predominant early adaptive event in our selection experiments and indicated that resistance-conferring DHFR mutations were likely to be selected only after PhoPQ signalling had been de-repressed.

The above experiments showed that mutations in *folA* and *mgrB* had a synergistic effect on the phenotype of trimethoprim-resistant *E. coli*. We had shown earlier that several mutations in DHFR, notably those at Trp30, are detrimental for its stability and result in aggregation, proteolysis and reduced levels [33]. Therefore, we hypothesized that enhanced expression due to PhoPQ de-repression could compensate for loss of mutant DHFRs due to mis-folding. To test this, we expressed wild type DHFR or its W30G/W30R mutant alleles from an IPTG-inducible promoter and checked the impact that expression level had on drug IC50 (Figure 3E). While trimethoprim IC50 was enhanced by ~3 fold upon overproduction of wild type DHFR, it was potentiated by x16- and x8-fold for W30G and W30R DHFR alleles respectively. We performed the same assay with W30F and W30L mutant alleles of DHFR. These alleles have comparable *in vivo* sensitivity to trimethoprim, but only W30L destabilizes the protein [33]. Overexpression affected the phenotype of W30F to a similar extent as wild type DHFR, while the IC50 of trimethoprim for *E. coli* expressing W30L DHFR was stimulated x8-fold upon overexpression (Figure 3E). Thus, mutant DHFR alleles resulted in higher gains in drug IC50 upon overexpression, explaining the synergy between *mgrB* and *folA* mutations, and this was related to their destabilizing effects on DHFR.

### Cost-compensation drives evolution of the PhoPQ transcriptional network during long-term antibiotic exposure

Having established a new mechanism for trimethoprim tolerance, and its effect as a potentiator of trimethoprim resistance, we next sought to understand the long-term evolutionary consequences of this adaptation. We established 3 lineages of *E. coli* evolving in high concentration of trimethoprim (300 ng/mL, x0.5-fold MIC) for ~350 generations (designated WTMP300 A, B and C). In agreement with results from short-term trimethoprim exposures (Figure 1A, Table 1), mutations in the *mgrB* gene or its promoter swept through all 3 lineages within the first ~20 generations of antibiotic exposure (Table S1). Subsequently, at this high drug pressure, there was rapid enrichment of resistant-bacteria that were eventually fixed in all 3 lineages (Figure 4A).

**Figure 4.**
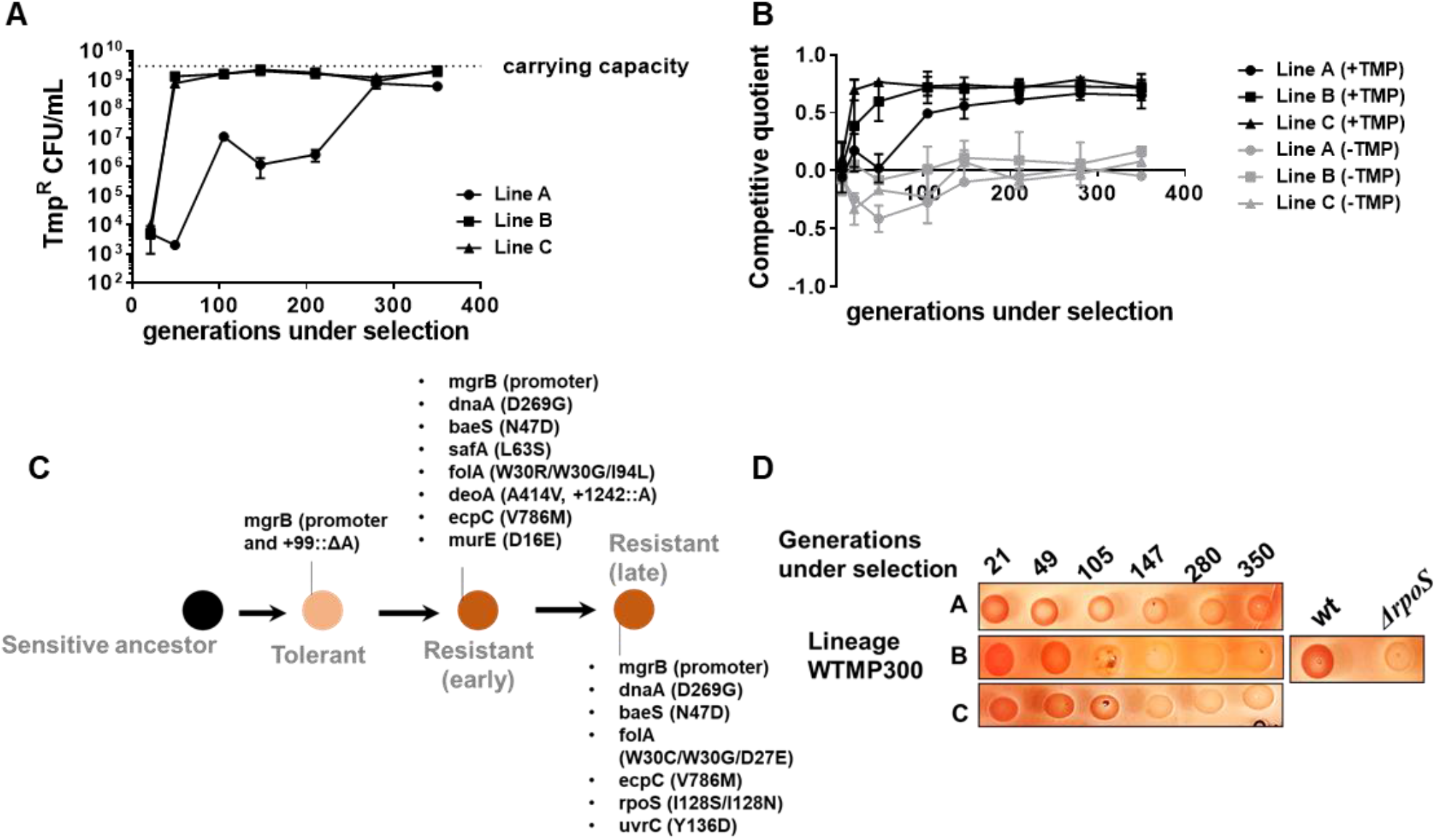
Genomic changes in *E. coli* adapting to high trimethoprim concentrations. **A**. Titre of trimethoprim-resistant bacteria in WTMP300 lineages A, B and C during the course of evolution. Each data point represents mean ± S.D. from 2-3 measurements. The typical carrying capacity of *E. coli* growing in drug-free medium (2-3 × 10^9^ CFU/mL) is indicated with a dotted line. **B**. Competitive quotient of WTMP300 lineages (A, B and C) in drug-free (gray) or trimethoprim supplemented (black) media were monitored by competition with *E. coli*-GFP (wild type *E. coli* expressing GFP). Higher values of competitive quotient indicate higher fitness relative to the wild type ancestor. Trimethoprim was used at 300 ng/mL. Each point represents mean ± S.D. from 2-3 measurements. **C.** Schematic representation of genomic changes associated with early and late adaptation to trimethoprim in WTMP300 lineage A (see Table S1 for complete list). **D**. Congo-red staining of WTMP300 lineages A, B and C to verify loss of active RpoS. Controls (wild type and *E. coli ΔrpoS*) are shown for reference. Representative data from 2-3 replicates are shown.

Next, we tracked adaptation of the evolving lineages to trimethoprim by competing them against *E. coli-*GFP. For all 3 lineages, competitive quotient rose steeply initially and then plateaued (Figure 4B). The initial rapid phase of adaptation was concordant with enrichment of trimethoprim-resistant bacteria (Figure 4A, B). Interestingly, IC50 of trimethoprim for resistant isolates from all three lineages plateaued early during evolution and did not change significantly over time (Figure S4). Thus, once resistance had evolved, its fixation in the population did not require further gains in drug MIC/IC50. On the other hand, competitive quotients of evolving populations in trimethoprim-free media initially decreased (Figure 4B), most likely due to the costs of mutations at the *mgrB* locus (Figure 1D). However, all three lineages recovered over time such that at 350 generations of evolution their relative fitness was similar to or slightly better than the ancestor (Figure 4B). Taken together, we interpreted these results to mean that trimethoprim-resistant bacteria enhanced their fitness over long-term evolution by amelioration of fitness costs, rather than enhancement in drug IC50.

To understand the genetic basis of these adaptations, we sequenced the genomes of 3 to 5 resistant isolates from Lineage A at three different timepoints i.e. 105, 147 and 350 generations of evolution (Table S1, Figure 4C). This allowed us to reconstruct the evolutionary genetic trajectory of trimethoprim-resistant bacteria in this lineage. Sequenced isolates from all three time points had mutations at the *mgrB* locus, re-iterating its role as a driver of early adaptation to trimethoprim (Table S1, Figure 4C). Mutations at the *folA* locus were also common among isolates at all 3 time points, explaining the high IC50 of trimethoprim for these isolates (Table S1, Figure 4C). In addition, some isolates harboured mutations at the *baeS* locus, a known regulator of efflux pump expression [43], and *safA*, a regulator of PhoPQ signalling [44, 45] (Table S1, Figure 4C). Strikingly, all sequenced isolates from 350 generations, but not earlier time points, harboured substitution of Ile128 to Ser or Asn in RpoS (Table S1, Figure 4C). The Ile128 residue of the RpoS sigma factor is critical for its activity, and hydrophilic substitutions at this site inactivate it [46]. In order to confirm loss of RpoS activity in these lineages, we exploited the differential staining of RpoS-deficient and RpoS-expressing *E. coli* by Congo Red. Congo Red stains curli fibres and cellulose, which are produced only by bacteria with active RpoS [47]. In agreement with results from genome-sequencing, all 3 lineages showed progressively poorer staining with Congo Red (Figure 4D). Thus, while resistance-associated genetic changes were observed early during evolution, loss-of-function mutations in *rpoS* were associated with adaptation during long-term exposure to trimethoprim.

Loss of RpoS, we had found, did not itself confer trimethoprim-resistance (Figure 1C). Further, since RpoS is stimulated by PhoPQ signalling, it was reasonable to expect that loss of RpoS activity compensated for the costs of *mgrB*-mutations. In support of this suggestion, RpoS-deficient bacteria were selected over RpoS-expressing bacteria only in a *ΔmgrB* background (Figure 5A). Moreover, this effect was relatively independent of the presence of trimethoprim (Figure 5A). RpoS activates the transcription of a number of stress-responsive and stationary phase genes. Several of these are up-regulated in an *mgrB*-knockout [38]. Could the costs of *mgrB* deletion be related to unrequired expression of stress and stationary phase genes? We tested this hypothesis by generating knockouts of 4 RpoS-regulated genes i.e. *gadX*, *cbpA*, *fic* and *sra* that are up-regulated in a *mgrB*-deficient background [38]. Knockouts of three of these, i.e. *gadX*, *cbpA* and *fic* were all selected over *E. coli ΔmgrB* independent of the presence of trimethoprim (Figure 5A). These data confirmed that loss of RpoS compensated for the fitness costs of PhoPQ de-repression by ameliorating the expression of unwanted stationary-phase and stress-related genes (Figure 5B).

**Figure 5.**
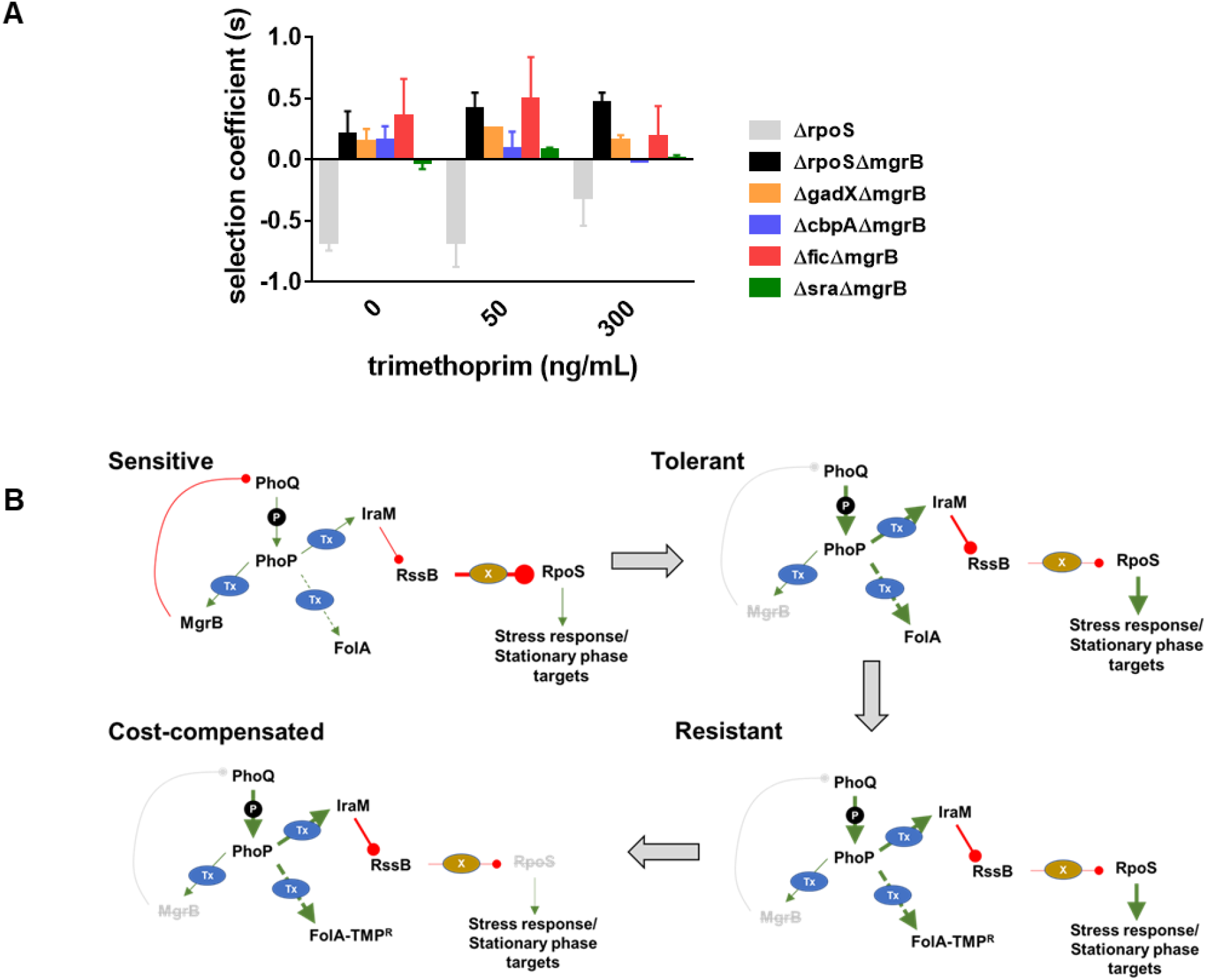
Compensation of fitness costs of *mgrB*-deficiency by loss of RpoS. **A.** Selection coefficient of *E. coli ΔrpoS* relative to wild type (gray bars), *E. coli ΔmgrBΔrpoS* relative to *E. coli ΔmgrB* (black bars) or indicated mutants in the *ΔmgrB* background relative to *E. coli ΔmgrB* (coloured bars). Appropriate strains were allowed to compete in the absence or presence of trimethoprim at the indicated concentrations for ~ 9 generations. Data represent mean ± S.E.M from 3 independent measurements. **B.** Schematic representation of adaptation of *E. coli* to high trimethoprim over long term evolution. Genetic changes and mechanisms associated with sensitive, tolerant, resistant and cost-compensated phenotypes are shown. The evolutionary sequence is indicated by the direction of arrows. Activating interactions are shown by green arrows, while inhibitory interactions are shown by red lines. Strength of each interaction is qualitatively represented by thickness of arrows. Indirect interactions are shown as discontinuous lines. Phosphorylation is indicated by ‘P’. Transcriptional changes are indicated by ‘Tx’. Proteolytic degradation is indicated by ‘X’. Inactivating mutations are represented by gray text with strikethrough.

### Strength of antibiotic selection determines the choice of adaptative strategy during the evolution of the PhoPQ-folA-RpoS transcriptional network

Our experiments so far identified two possible strategies available to *E. coli* for enhancing fitness subsequent to evolving tolerance, i.e. resistance and cost-compensation. We next asked what motivated the ‘choice’ of strategy that an evolving bacterial population would adopt. Cost-compensation is known to allow resistant bacteria to enhance their fitness in drug-free or low antibiotic conditions. Therefore, we argued that antibiotic concentration may determine which adaptive strategy was favoured. In order to test this, we mixed tolerant (*E. coli ΔmgrB*), cost-compensated tolerant (*E. coli ΔmgrBΔrpoS*) and resistant (TMPR1) strains (initial mixing ratio 98:1:1) and allowed them to compete for ~9 generations in different concentrations of trimethoprim. In the absence of trimethoprim or in low concentrations of the antibiotic, *ΔmgrBΔrpoS* was strongly enriched over *ΔmgrB* and TMPR1 (Figure 6A). However, at higher drug concentrations, TMPR1 was favoured over its competitors (Figure 6A). This clear dependency on trimethoprim concentration suggested that at low drug pressures, evolutionary outcomes were likely to be biased in favour of cost-compensation rather than resistance-evolution.

**Figure 6.**
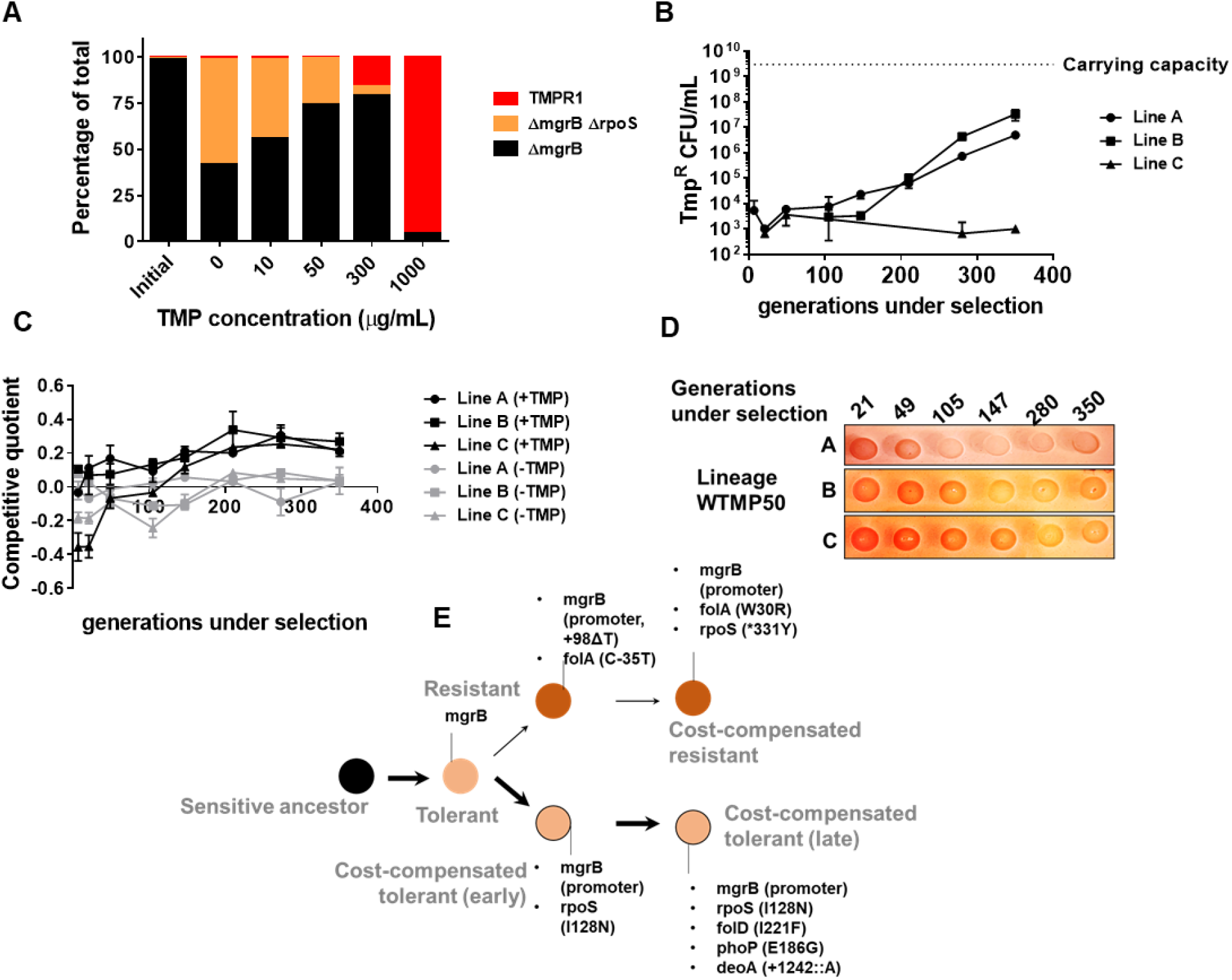
Genomic changes in *E. coli* adapting to low trimethoprim concentrations. **A**. Three-strain competition between *E. coli ΔmgrB* (tolerant), *E. coli ΔmgrBΔrpoS* (cost-compensated tolerant) and *E. coli* TMPR1 (resistant). Initial mixing ratio of strains was 98:1:1 respectively. Ratios of strains after ~ 9 generations of competition at the indicated concentration of trimethoprim are shown. Data are representative of 3 replicates. **B**. Titre of trimethoprim-resistant bacteria in WTMP50 lineages A, B and C during the course of evolution. Each data point represents mean ± S.D. from 2-3 measurements. The typical carrying capacity of *E. coli* growing in drug-free medium (2-3 × 10^9^ CFU/mL) is indicated with a dotted line. **C**. Competitive quotient of WTMP50 lineages A, B and C in drug-free (gray) or trimethoprim supplemented (black) media were monitored by competition with *E. coli*-GFP (wild type *E. coli* expressing GFP). Higher values of competitive quotient indicate higher fitness relative to the wild type ancestor. Trimethoprim was used at 50 ng/mL. Each point represents mean ± S.D. from 2-3 measurements. **D**. Congo-red staining of WTMP50 lineages A, B and C to verify loss of active RpoS in all lines. Representative data from 2-3 replicates are shown. **E**. Schematic representation of genomic changes associated with early and late adaptation to trimethoprim in WTMP50 lineage A (see Table S2 for complete list). Two parallel lineages branching out from tolerant bacteria in the WTMP50 lines are shown.

We empirically tested the above prediction in 3 replicate lineages of *E. coli* K-12 MG1655 evolving under low trimethoprim pressure (50 ng/mL, 0.08x MIC, designated WTMP50 A, B and C). In support of our hypothesis, resistant bacteria did not exceed more than 1% of the population in these lineages for the duration of the experiment (Figure 6B). Indeed, in WTMP50 Lineage C the fraction of resistant bacteria stayed below 0.001% of the total population throughout the experiment (Figure 6B). Gain in relative fitness of WTMP50 lineages over the ancestor was lower than WTMP300, and showed very poor concordance with enrichment of trimethoprim-resistant bacteria over time (Figure 6C). Importantly, like WTMP300, all 3 WTMP50 lineages got fitter in antibiotic-free medium over time (Figure 6C) and lost RpoS activity (Figure 6D). These data suggested that there were at least 2 parallel evolving adaptive trajectories in the WTMP50 lines, i.e. resistant bacteria, that made up a minority of the population and cost-compensated, tolerant bacteria that made up the majority of the population. Genome sequencing of resistant and tolerant isolates from 147 and 350 generations under selection from WTMP50 Lineage A, confirmed that this was indeed the case (Table S2, Figure 6E). The genetic trajectory of resistant bacteria in WTMP50 was very similar to the WTMP300, in that mutations in *folA* were acquired before cost-compensatory mutations in *rpoS* (Table S2, Figure 6E). On the other hand, tolerant bacteria acquired *rpoS* mutations earlier during evolution and did not acquire mutations in folA even after long term antibiotic exposure (Table S2, Figure 6E). Thus, cost-compensation was indeed the preferred strategy for adaptation at lower antibiotic concentrations.

Drug concentrations are generally equated with strength of selection for antibiotic resistance [48, 49]. However, selection strength is determined by intrinsic factors as well such as the genetic background [40, 50]. Therefore, we tested whether the link between selection strength and evolutionary trajectory would hold true in another genetic background. We have shown earlier that Lon protease-deficient *E. coli* are trimethoprim-tolerant due to a *folA*-independent mechanism [50]. For *E. coli Δlon* it is expected, then, that 300 ng/mL of trimethoprim would represent a milder selection strength than wild type. Therefore, we established 3 lineages of *E. coli Δlon* that experienced trimethoprim (300 ng/mL, 0.5× MIC, designated LTMP300 A, B and C) for 350 generations and examined their adaptation over time. At the phenotypic level, LTMP300 lineages resembled WTMP300 in terms of fixation of resistant-bacteria (Figure 7A). However, at the genotypic level there was substantial similarity between LTMP300 and WTMP50 lineages. We found no *folA* mutations in resistant isolates from the LTMP300 A lineage (Table S3, Figure 7B). Instead, trimethoprim-resistant bacteria derived from an early time point (147 generations) had already accumulated inactivating mutations in *rpoS* in addition to mutations in *mgrB* (Table S3, Figure 7B). Thus, while the LTMP300 lineages had the genetic signature of evolution at low selection strength, they phenotypically resembled high selection strength. This discrepancy was explained by the fact that *Δlon* and *ΔmgrB* showed an additive effect on trimethoprim IC50 (Figure 7C). While loss of *mgrB* alone resulted in drug-tolerance, together with Lon-deficiency it resulted in drug-resistance. Importantly, once *mgrB* mutations were acquired, LTMP300 lineages evolved by compensatory mutations rather than resistance-conferring mutations in *folA*, similar to WTMP50. These results, therefore, re-iterated the role of selection strength in determining the genetic trajectory followed by *E. coli* during adaptation to trimethoprim (Figure 7D).

**Figure 7.**
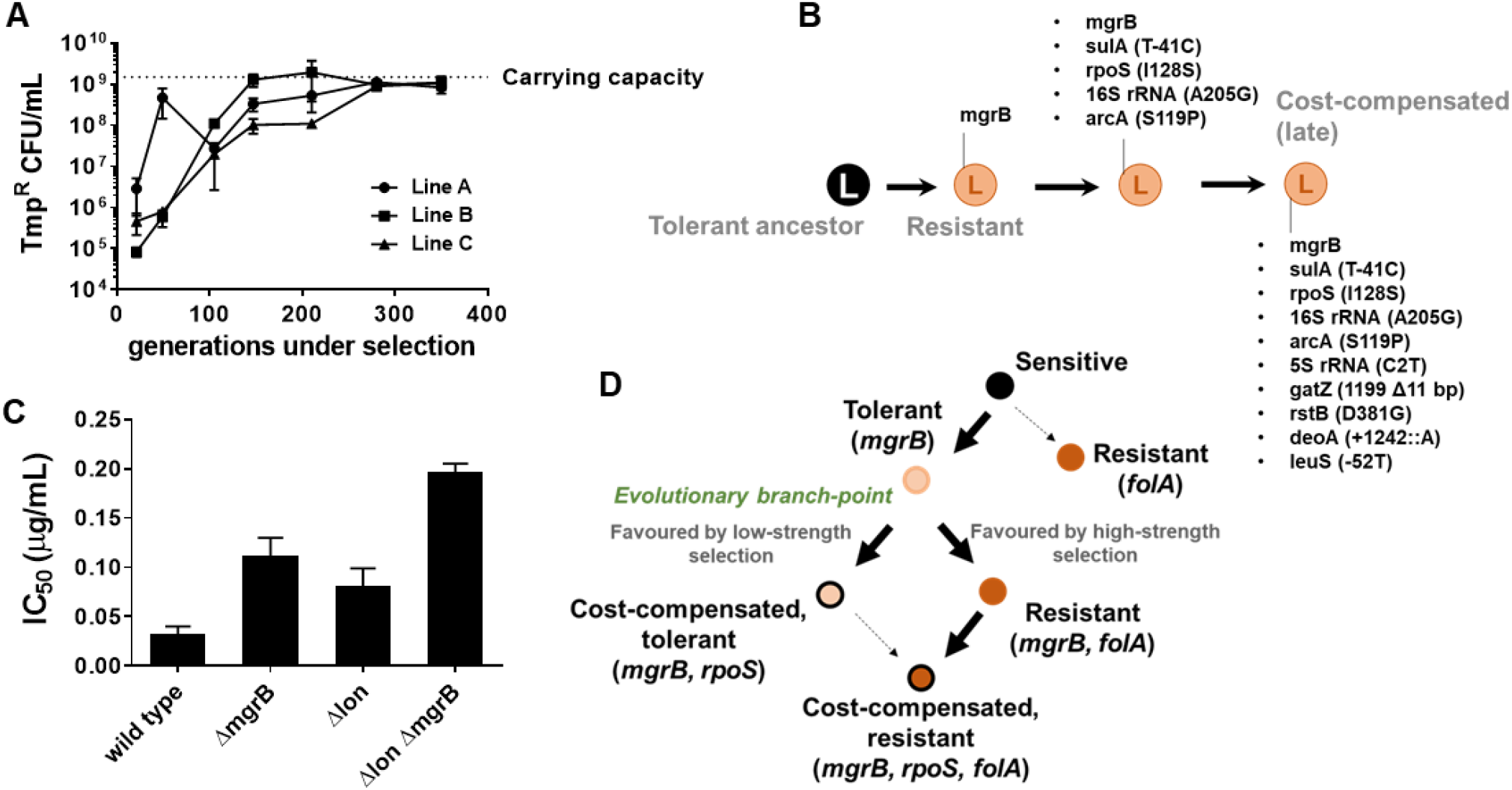
Genomic changes in *E. coli Δlon* adapting to high trimethoprim concentrations. **A**. Titre of trimethoprim-resistant bacteria in LTMP300 lineages A, B and C during the course of evolution. Each data point represents mean ± S.D. from 2-3 measurements. The typical carrying capacity of *E. coli Δlon* growing in drug-free medium (1-2 × 10^9^ CFU/mL) is indicated with a dotted line. **B**. Schematic representation of genomic changes associated with early and late adaptation to trimethoprim in LTMP300 lineage A (see Table S3 for complete list). **C**. Additive effect of *Δlon* and *ΔmgrB* mutations on IC50 of trimethoprim for *E. coli*. Mean ± S.D. values of IC50 from 3 independent experiments are plotted. **D**. Model of adaptation in the PhoPQ-folA-RpoS axis showing dependency of genetic changes and associated phenotypes on drug pressure. Implicated genetic loci are indicated in parentheses.Evolutionary transitions identified in this study are shown as solid black arrows. Possible transitions that are unlikely to occur based on the results from this study, are shown as dotted arrows.

## Discussion

The PhoPQ signalling cascade is a well-studied two-component system, not just in Enterobacteria but also other species. Traditionally thought to be regulated by Mg^2+^ [51], this system also responds to low pH [44] and periplasmic redox state [52]. Mounting evidence in recent years has thrown light on its role in resistance to antibiotics, particularly antimicrobial peptides such as polymyxins [53]. The clinical relevance of these findings was brought to the fore when a number of studies identified inactivating mutations in *mgrB* in carbapenem/colistin-resistant *Klebsiella pneumoniae* [54–56]. De-repression of PhoPQ in *Klebsiella* results in alterations in the structure of Lipid A, conferring resistance to colistin [57]. Inactivation of *mgrB* also collaterally enhances virulence in *Klebsiella* [57], a particularly worrying observation given that polymyxins represent the last line of defence against hospital-borne Gram-negative infections at the moment. Similarly in *E. coli*, C18G, another antimicrobial peptide, leads to growth arrest and filamentation by activating PhoPQ signalling [21].

In the context of trimethoprim resistance, mutations in *mgrB* were reported earlier by Baym et al. (2016) from drug-resistant *E. coli* evolving on mega petri-plates [58], though a mechanistic basis for this change was not known. Transcriptional up-regulation of *folA* upon loss of *mgrB*, a main result from our study, fills this lacuna. Additionally, by showing the synergistic interaction between mutations in *mgrB* and *folA*, our study also establishes how loss of *mgrB* facilitates resistance evolution. These results are in line with other studies that have identified the facilitatory role that drug tolerance plays in the evolution of resistance [59]. Early inactivation of *mgrB* in *E. coli* upon trimethoprim exposure raises two questions. Firstly, how does PhoPQ enhance the expression of *folA?* The only known transcriptional regulator of *folA* in *E. coli* is the TyrR transcription factor [60]. However, we were unable to detect any change in expression level of DHFR protein in a *tyrR* deletion strain (Figure S5). Given the wide-spread gene regulatory impact of PhoPQ de-repression in *E. coli*, including its effects on other transcription factors, its effects on *folA* levels are likely to be secondary or tertiary. Importantly, these indirect regulatory effects of PhoPQ de-repression were sufficient to drive the selection of *mgrB* mutations. Further investigation into the mechanism of *folA* promoter up-regulation in *mgrB*-deficient *E. coli* is likely to reveal additional regulators for this house-keeping gene.

Secondly, is inactivation of MgrB relevant in clinical or environmental strains of *E. coli*? We detected 4 kinds of mutations at the *mgrB* locus in our experiments, the most frequent being disruption of the promoter of *mgrB*. Additionally, we found single nucleotide insertion or deletions within the coding region. These mutations are expected to result in a longer or shorter protein with markedly altered C-terminal sequence as a result of translational frame-shift (Figure S6). We searched for protein sequences similar to these frame-shifted alleles of MgrB in the non-redundant protein sequence database. BLAST analysis revealed 9 clinical and environmental *E. coli* strains from 4 different pathogen surveillance and sequencing projects that had altered C-terminal sequences similar to the mutant alleles found by us (Figure S6). Indeed, one of these strains, *E. coli* NCTC9075 from the collection of Public Health England, harboured exactly the same deletion in *mgrB* (+100 ΔA) that we identified in some of our laboratory-evolved strains (Figure S6). Additionally, we also identified mutant MgrB sequences from 3 isolates of *Salmonella enterica* and 1 isolate of *Citrobacter freundii* (Figure S7). Thus, the relevance of MgrB in trimethoprim tolerance, or indeed in other phenotypes of pathogenic Enterobacteria, are likely to be far more significant that currently understood.

Though we have focussed on the compensatory effects of RpoS mutations in this study, genomics analyses revealed a number of other loci that accumulated mutations in our laboratory evolution experiments. For instance, mutations in the *deoA* locus, predicted to cause pre-mature termination of the DeoA protein, were detected in resistant isolates from all selection schemes employed by us (Table 1, S1, S2, S3). DeoA, a thymidine phosphorylase, is part of the pyrimidine catabolism pathway in *E. coli* [61]. Deactivation of this pathway is known to be beneficial for *E. coli* harbouring hypomorphic DHFR alleles by reducing the demand for *de novo* pyrimidine biosynthesis [62]. Interestingly, mutations in some of the other loci from our study showed strict dependency on drug concentration used during selection. Most notably, mutations in *baeS*, *ecpC* and *dnaA* genes, which code for two component kinase, fimbrial chaperone and replication initiator proteins respectively, were found in a majority of resistant isolates from the WTMP300 lineages, but not in WTMP50 lineages (Table S1, S2). BaeS, sensor kinase of the BaeRS two-component system, has been associated with multi-drug resistance in a number of bacteria including *E. coli* due to its effect on the expression of efflux pumps [43]. The fimbrial chaperone EcpC, on the other hand, has never been associated with resistance to antibiotics, and is potentially a novel locus worth investigating. Point mutations in DnaA (D269G and D280G) that were detected in trimethoprim-resistant isolates from WTMP300 lineages are expected to compromise its ATP-binding [63]. Indeed, D269 lies in the ‘Sensor I’ motif of DnaA and its mutation to Ala reduces ATPase activity of the protein *in vitro* [64]. We envisage two possible reasons for these mutations in DnaA to be selected during adaptation to trimethoprim. Firstly, thymine-less death, induced by thymine starvation or trimethoprim treatment, can be prevented in *E. coli* by lowered rates of replication initiation [65]. Mutations in DnaA that compromise its ATPase activity would help achieve this. Secondly, DnaA mutations in clinical and laboratory strains of *Mycobacterium tuberculosis* have been associated with isoniazid resistance due to altered gene expression profiles [66], and a similar mechanism may be operative in the case of *E. coli* and trimethoprim. The appearance of mutations at these loci only at high antibiotic pressure are likely to be a result of the higher pressure for resistance.

We also observed mutations specific to genetic background. Most notable of these were mutations in 5S and 16S ribosomal RNAs, promoter of *sulA* and the ArcB transcriptional regulator in trimethoprim-resistant isolates evolved from the *Δlon* strain. The Lon protease degrades a number of house-keeping proteins, but can also degrade misfolded mutant proteins [67]. We have shown in the past that a Lon-deficient background permits access to a larger repertoire of mutations to adapting bacterial populations [50]. This phenomenon may very well explain why certain mutations were detected only in the Lon-deficient background. Further, mutations in the *sulA* promoter are expected to prevent its expression [68], thus alleviating the accumulation of this cell division inhibitor in the Lon mutant. Loss of *sulA* is advantageous for *E. coli Δlon* as it rescues the hyper-filamentation defect of this strain [50, 69], which may explain why mutations in the *sulA* promoter were identified only in the LTMPR300 lineages.

To our knowledge, the present study is one of a kind in being able to trace the evolutionary trajectory of a gene regulatory network from its initial perturbation in response to selection, to eventual evolutionary rescue by compensatory mutations. Three main observations from this study significantly contribute to a general understanding of how gene regulatory networks change in response to selection. First, changes in gene regulatory networks can alter the fitness landscape of adaptive mutations. The evolutionary potential of gene regulatory changes is well-recognized particularly in the context of vertebrate development [70]. Similarly, gene regulatory changes as mediators of evolution has been suggested in a series of synthetic biology experiments in prokaryotes [71]. Our study demonstrates that gene regulatory mutations can also act as facilitators of subsequent evolutionary change by shifting the fitness landscape of mutant alleles. This finding adds a new dimension to understanding the role of gene regulatory networks in mediating evolution of new functions. Further, mutant alleles of enzymes, like in the case of DHFR, are known to often display trade-offs with stability [33, 72]. The role of gene regulatory networks in negotiating these trade-offs needs to be factored into understanding the genetics of evolutionary adaptation.

Secondly, cost-compensation emerged as an important force driving the evolution of the PhoPQ-folA-RpoS network. Evolution of gene regulatory networks is thought to be primarily under stabilizing selection, in light of empirically observed properties such as developmental canalization and mutational robustness [73]. Compensatory evolution in regulatory networks have been understood mainly in the context of cis or trans-regulatory changes that minimize variation in gene expression [74, 75]. Our results demonstrate that the fitness effects of adaptive mutations can be significant. As a result, compensation of fitness cost can also drive additional changes in regulatory networks. Importantly, loss of RpoS activity, a key cost-compensatory change in our experiments, is a commonly encountered trait in Enterobacteria [76–78]. RpoS mutants are associated with fitness advantage in aging bacterial colonies [79], and natural isolates of *E. coli* often have inactive alleles of this sigma factor [76–78, 80, 81]. The driving force behind loss of RpoS in Enterobacteria is thought to be selection for enhanced growth rate [76, 81]. Since RpoS competes with house-keeping sigma factor RpoD for RNA polymerase occupancy, loss of RpoS drives growth and replication, and compromises stress tolerance as a trade-off [76, 81]. The results of our study support this thesis. However, in our study, selection of RpoS mutants was driven in response to gene regulatory perturbation and not by environmental factors. Given the high prevalence of RpoS-mutations in natural isolates of Enterobacteria, it is very likely that they may serve as compensatory changes for other gene regulatory perturbations as well.

Finally, the dependency of evolutionary strategy on selection pressure is a key finding from this study (Figure 6D). Over the last decade there has been significant attention given to how sub-lethal drug concentrations, such as those found in natural reservoirs of bacteria, alter the evolution of drug resistance [40–42, 48, 82–84]. The present study provides direct evidence linking drug concentration to the genetics of adaptation to antibiotics in bacteria. We also provide a mechanistic understanding for this link. Cost-compensation by second-site mutations is well-known for antimicrobial resistance and is indeed a widely-accepted phenomenon [85–89]. It is thought to occur under relaxed selection, i.e. when drug is not present in the environment. Our study demonstrates that compensation of cost and resistance evolution can also be thought of alternative strategies for enhancing fitness in the presence of the antibiotic. The relative gain in fitness from each of these strategies is determined not just by drug concentration but also by genetic background, and both these factors have vital influence on the evolutionary outcomes of bacteria under antibiotic selection.

## Materials and Methods

### Strains, plasmids and culture conditions

*E. coli* K-12 MG1655 or its derivates were cultured in Luria-Bertani broth supplemented with appropriate antibiotics at 37 °C with shaking (180-200 rpm). The various strains and plasmids used in this study and their sources are listed is Table 2. Single gene knockouts were obtained from the Keio Collection [90, 91](from the NBRP Resource, National Institute of Genetics, Japan) and moved into the MG1655 background using P1 transduction. For double gene knockouts, antibiotic marker cassette was first removed using Flp recombinase (pCP20-flp plasmid) [92], and then subjected to P1 transduction.

**Table 2.**
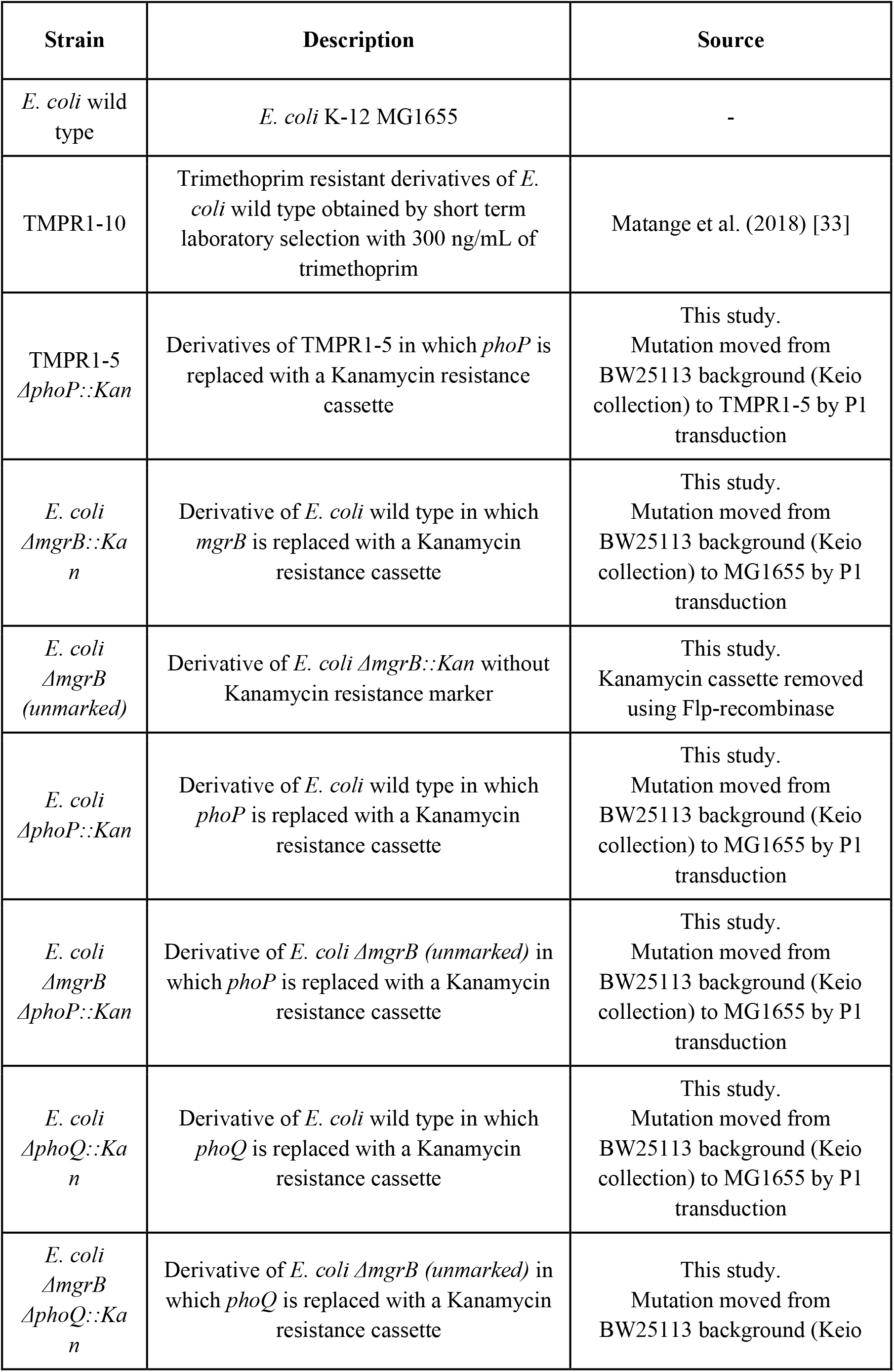
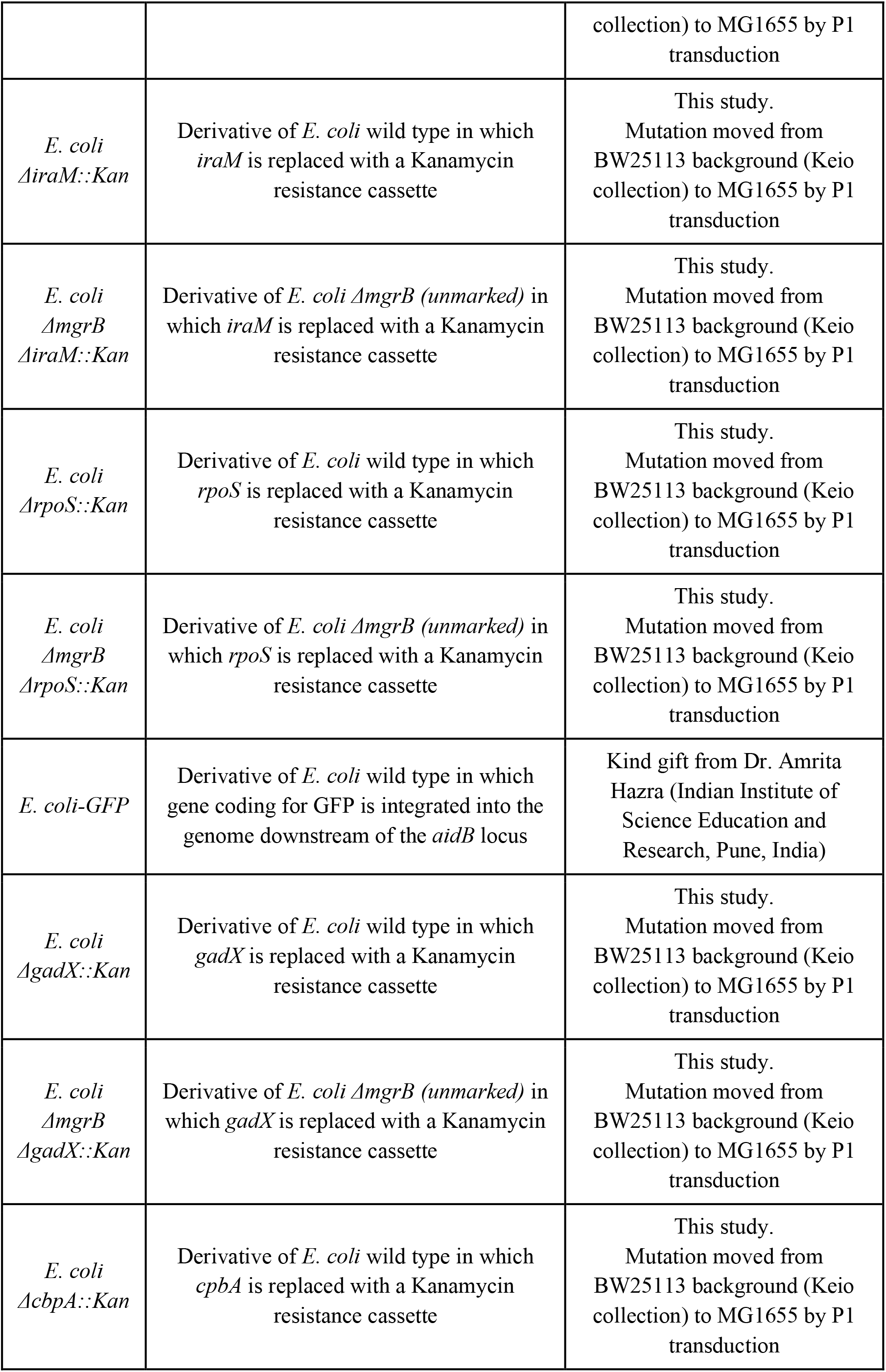
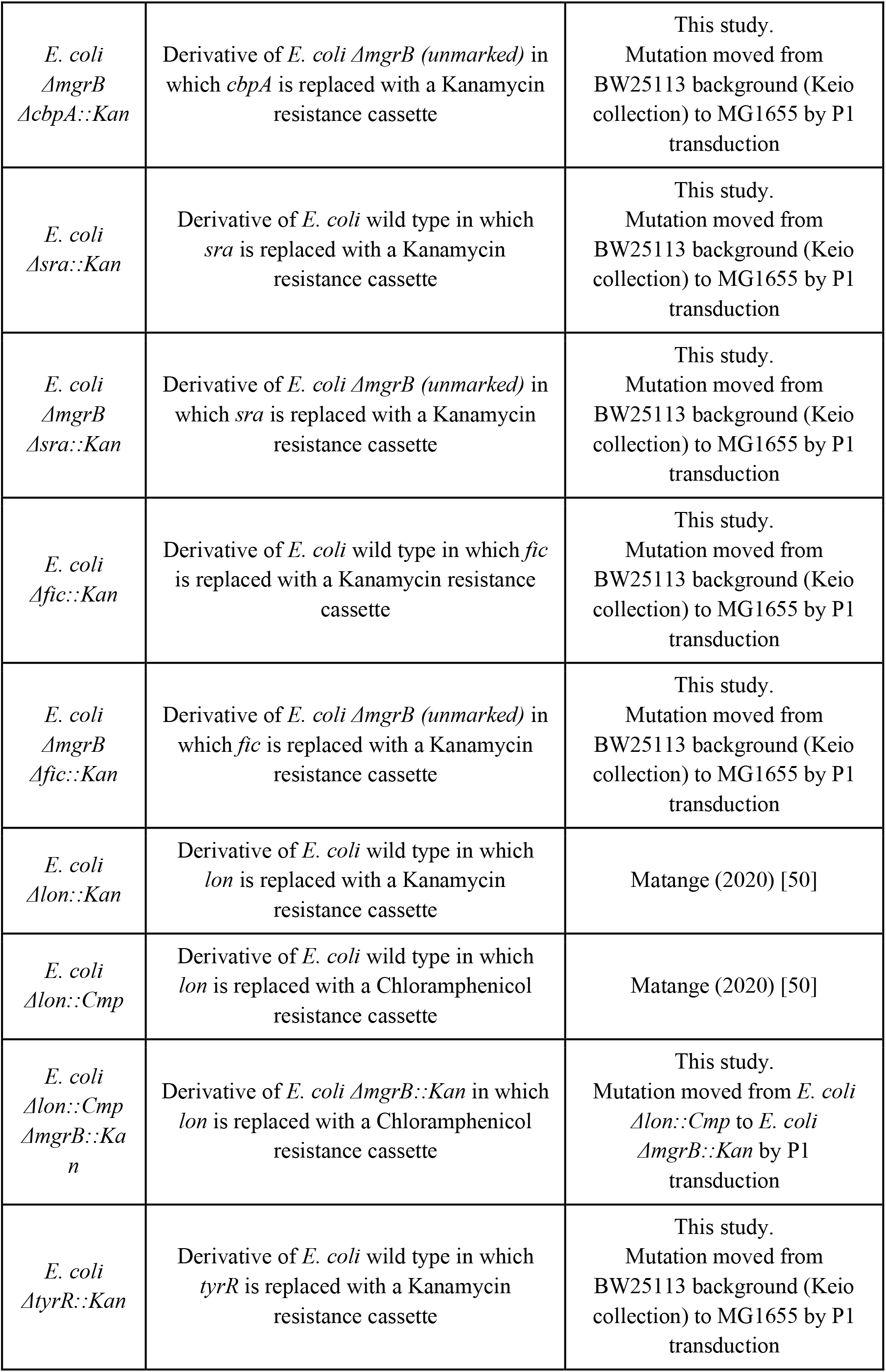
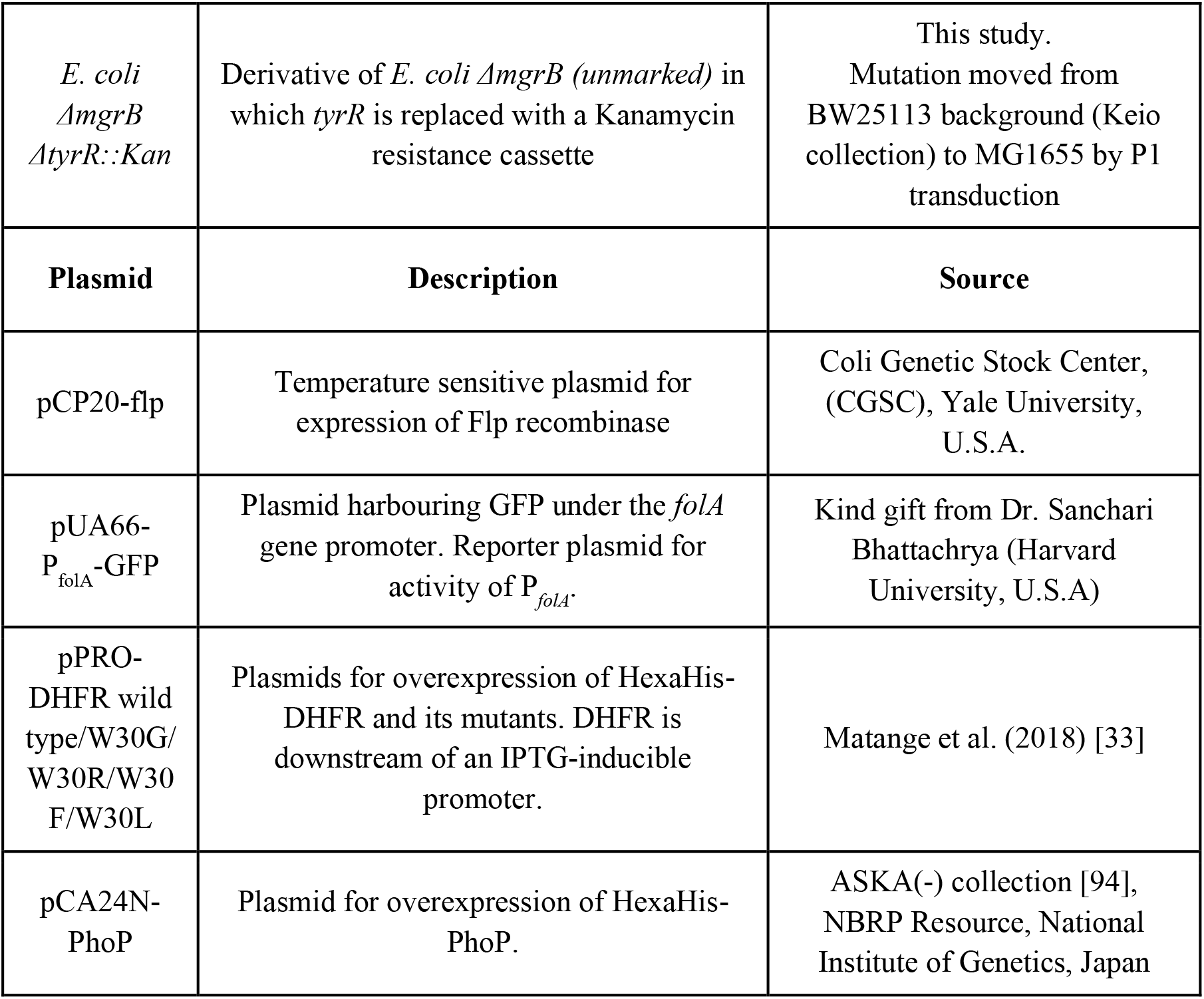
List of strains and plasmids used in this study.

Antibiotics used in this study were purchased from Himedia (India) or Sigma-Merck (U.S.A., Germany). Kanamycin (30 μg/mL), chloramphenicol (25 μg/mL) or ampicillin (100 μg/mL) were added to liquid or semi-solid media just before inoculation as required. For isolating trimethoprim-resistant mutants from laboratory evolution experiments Luria-Bertani agar was supplemented with 1 μg/mL trimethoprim.

### Antibiotic sensitivity

Trimethoprim sensitivity of gene knockouts or evolved clones was tested using the following methods:

#### Minimum Inhibitory Concentration (MIC)

MIC determination was performed using E-test. Trimethoprim strips (x – x μg/mL) were procured from Himedia (India) and used as per manufacturer’s instruction.

#### Inhibitory Concentration-50 (IC50)

IC50 of trimethoprim was determined using a broth dilution assay as described in Matange et al. (2019) [40]and Matange et a. (2018) [33].

#### Colony formation

Colony forming efficiency of wild type or mutant *E. coli* at was calculated by spotting 10 μl of neat or serially diluted (10-fold dilution series) late log-phase cultures on Luria-Bertani agar supplemented with 0, 50, 100, 300, 500 and 1000 ng/mL trimethoprim. Plates were incubated for 18h at 37 °C before counting colonies.

#### Colonization efficiency

Wild type or mutant *E. coli* cultures were grown to saturation and then serially diluted. Increasing bacterial CFUs (~10^4^, 10^5^,10^6^, 10^7^) were added to 1 mL of Luria-Bertani broth supplemented with 0, 500, 1000 or 5000 ng/mL trimethoprim. Cultures were grown for 18h at 37 °C and optical density (at 600 nm) was measured in a spectrophotometer.

### Relative fitness and competitive growth assays

Relative fitness of gene knockouts or evolved clones were estimated using competition assays. Test strains were mixed with ancestral *E. coli* from saturated mono-cultures (1:1, unless otherwise mentioned) in 1-3 mL Luria-Bertani broth supplemented with trimethoprim at the appropriate concentration. Strains were allowed to compete for ~ 9 generations.

For 2-strain or 3-strain competitions, CFU/mL for each strain were determined by plating serially diluted cultures on antibiotic-free (total CFU) and kanamycin (knock-out strains) or trimethoprim (trimethoprim resistant isolates) supplemented Luria-Bertani agar. Relative fitness (w) or selection coefficient (s) were calculated using the following formulae:

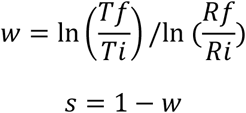

where, Tf and Ti are the final and initial CFU/mL of the test strain and Rf and Ri are the final and initial CFU/mL of the reference strain respectively.

For competitive quotients, GFP-expressing derivative of *E. coli* wild type (*E. coli*-GFP) was used as the reference strain. At the end of the competition experiment, 200 μL of each mixed culture was aliquoted into a 96-well plate and GFP fluorescence and optical density (at 600 nm) was estimated in a multi-mode plate reader (Varioskan-Thermo Scientific or Enlight-Perkin Elmer). For each individual experiment 2 controls were maintained, i.e. *E. coli-GFP* alone and control competition between *E. coli-*GFP and wild type. GFP fluorescence was normalized to optical density (nGFP) and used to calculate competitive quotient using the following formula:

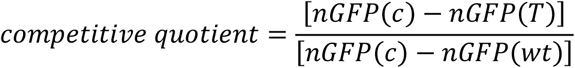

where, nGFP(c), nGFP(T) and nGFP(wt) are the normalized GFP fluorescence values of *E. coli*-GFP alone, mixed culture of *E. coli*-GFP and test strain, and mixed culture of *E. coli*-GFP and wild type respectively.

### Immunoblotting

Immunoblotting was used to determine the expression level of DHFR in mutants or resistant isolates as described in Matange (2020) [50]. FtsZ was used as loading control. Anti-DHFR IgG was used at a concentration of 100 ng/mL. Anti-FtsZ (kind gift from Prof. Manjula Reddy, CCMB, India) polyclonal antiserum was used at a dilution of 1:50,000.

### Activity of *folA* gene promoter

The activity of the promoter of *folA* was measured in wild type or mutant *E. coli* using the pUA66-P_folA_-GFP reporter plasmid (kind gift from Dr. Sanchari Bhattacharya, CCCCC). Plasmid reporter was transformed into appropriate strains and GFP fluorescence was measured over growth in a multi-mode plate reader (Varioskan-Thermo Scientific). For comparison across strains GFP fluorescence was normalized to optical density.

### Purification of His-tagged PhoP

Hexa-His tagged PhoP was overexpressed in *E. coli* K-12 MG1655 and purified using Ni-NTA affinity chromatography. Bacterial strain harbouring pCA24N-PhoP plasmid was grown in 50-100 mL Luria-Bertani broth till an optical density of ~0.8. Protein production was induced using IPTG (500 μM). Induction was carried out for 3 hours at 37 °C, following which bacterial cultures were centrifuged at 3000g for 10 mins. Bacterial pellets were stored at -80 °C until needed. Cells were lysed by sonication in Lysis Buffer (Tris-HCl pH 8.0 100 mM, NaCl 100 mM, β-mercaptoethanol 1 mM, glycerol 10%) and loaded onto a Ni-NTA-agarose resin (Bio-Rad, U.S.A). The resin was washed using Wash buffer (Tris-HCl pH 8.0 100 mM, NaCl 300 mM, β-mercaptoethanol 1 mM, imidazole 20 mM) and eluted using Elution buffer (Tris-HCl pH 8.0 100 mM, NaCl 100 mM, β-mercaptoethanol 1 mM, imidazole 300 mM, glycerol 10%). Eluted protein was desalted using PD-10 desalting columns (Sigma-Merck, U.S.A/Germany). Concentration of the desalted protein was estimated using Bradford’s method and the protein was stored at -80 °C until needed.

### Electrophoretic mobility shift assay (EMSA)

For EMSA, biotinylated probes shown in Figure S3 were synthesized (Sigma, U.S.A) and annealed to a final concentration of 25 pmol/μL. They were further diluted to 25 fmol/μL in nuclease-free water. Recombinant purified PhoP protein was phosphorylated in vitro by incubating with 20 mM acetyl-phosphate and 10 mM MgCl_2_ for 45 mins at 37 °C. The phosphorylated protein was used immediately for further experiments. For binding and visualization LightShift Chemilunescent EMSA kit (Thermo, U.S.A) was used as per manufacturer’s instructions.

### Laboratory evolution of trimethoprim resistance

For long-term laboratory evolution of trimethoprim resistance, replicate lineages (3 for each drug concentration/genotype) were established using serial passaging in trimethoprim-supplemented Luria-Bertani broth (2 mL). Bacteria (1 %) were transferred to fresh media every 10-14 hours such that each growth cycle had between 6 and 7 generations. Laboratory evolution was continued for ~350 generations (i.e. 50 growth cycles). Periodically an aliquot of culture from each lineage was mixed with an equal volume of sterile glycerol (50%) and frozen at -80 °C for analysis.

The titre of trimethoprim-resistant bacteria from each of the frozen stocks was determined by spotting 10 μL of serially-diluted culture on Luria-Bertani agar supplemented with trimethoprim (1 μg/mL). Resistant colonies (3-6) were selected at random for MIC determination or sequencing. Isolation of tolerant bacteria from WTMP50 lineages was done by spotting serially diluted cultures on drug-free medium. Tolerant isolates (3-6) were re-inoculated and their MIC was determined to confirm that they were not resistant to trimethoprim.

### Congo Red staining of bacterial colonies

Congo Red staining was used to determine RpoS activity in evolved lineages. Saturated cultures (10 μl) of appropriate strains were spotted on petri-plates containing YESCA medium (0.1% w/v yeast extract, 1% w/v casein hydrolysate, 2% w/v agar) supplemented with Congo red (50 μg/mL) and Coomassie Brilliant Blue G-250 (10 μg/mL). Plates were incubated at 28 °C to induce Curli fibres and photographed after ~24h of growth.

### Sequencing and genome analysis

Genomic DNA was extracted from evolved resistant/tolerant isolates using phenol:chloroform:isoamyl alcohol extraction as described in Matange et al. (2019) [40]. Extracted gDNA was then cleaned up using QiaAMP DNA purification spin columns (Qiagen, Germany) and quantitated spectrophotometrically. Integrity of the prepared gDNA was verified by gel electrophoresis before further applications.

The *folA* locus (encompassing the coding region and 195 bp upstream promoter) was PCR amplified from gDNA of trimethoprim-resistant isolates as described in Matange et al. (2018) [33] and Sanger sequenced.

For whole genome resequencing, paired-end whole-genome Next-Generation Sequencing (NGS) was performed on a MiSeq system (Illumina, USA) with read lengths of 150 bps. Processed reads were aligned to the reference genome *E. coli* K-12 MG1655 (NZ_CP025268) using bowtie2. A cut-off of a minimum of 20x coverage was employed, which made >97% of the reads for each of the samples permissible for analysis. Variant calling and prediction of new junctions in the genome to identify large structural mutations such as insertions and deletions was done using Breseq using default settings [93]. All variants and new junctions that were already present in the wild type ancestor were excluded.

Sanger sequencing services were provided by FirstBase (Malaysia). Library preparation and NGS services were provided by Bencos (India).

## Acknowledgements

The authors would like to acknowledge Dr. Amrita Hazra (IISER, Pune, India) and Dr. Manjula Reddy (CCMB, Hyderabad, India) for strains and reagents. The authors would like to acknowledge National Bioresource, NIG, Japan, for *E. coli* mutants. This project was funded by the INSPIRE program (Department of Science and Technology, Govt. of India) and IISER-Pune. NM is a recipient of the SERB-Research Scientist Award (Science Education and Research Board, Govt. of India).

## References

1. Hill, M.S., P. Vande Zande, and P.J. Wittkopp, Molecular and evolutionary processes generating variation in gene expression. Nat Rev Genet, 2021. 22(4): p. 203–215.

2. Kemble, H., P. Nghe, and O. Tenaillon, Recent insights into the genotype-phenotype relationship from massively parallel genetic assays. Evol Appl, 2019. 12(9): p. 1721–1742.

3. Kim, Y.A. and T.M. Przytycka, Bridging the Gap between Genotype and Phenotype via Network Approaches. Front Genet, 2012. 3: p. 227.

4. Levine, M. and E.H. Davidson, Gene regulatory networks for development. Proc Natl Acad Sci U S A, 2005. 102(14): p. 4936–42.

5. Singh, A.J., S.A. Ramsey, T.M. Filtz, and C. Kioussi, Differential gene regulatory networks in development and disease. Cell Mol Life Sci, 2018. 75(6): p. 1013–1025.

6. Shen-Orr, S.S., R. Milo, S. Mangan, and U. Alon, Network motifs in the transcriptional regulation network of Escherichia coli. Nat Genet, 2002. 31(1): p. 64–8.

7. Perraud, A.L., V. Weiss, and R. Gross, Signalling pathways in two-component phosphorelay systems. Trends Microbiol, 1999. 7(3): p. 115–20.

8. Mitrophanov, A.Y. and E.A. Groisman, Signal integration in bacterial two-component regulatory systems. Genes Dev, 2008. 22(19): p. 2601–11.

9. Valverde, C. and D. Haas, Small RNAs controlled by two-component systems. Adv Exp Med Biol, 2008. 631: p. 54–79.

10. Hengge, R., The two-component network and the general stress sigma factor RpoS (sigma S) in Escherichia coli. Adv Exp Med Biol, 2008. 631: p. 40–53.

11. Stephenson, K. and R.J. Lewis, Molecular insights into the initiation of sporulation in Gram-positive bacteria: new technologies for an old phenomenon. FEMS Microbiol Rev, 2005. 29(2): p. 281–301.

12. Beier, D. and R. Gross, Regulation of bacterial virulence by two-component systems. Curr Opin Microbiol, 2006. 9(2): p. 143–52.

13. Sivaramakrishnan, S. and P.R. de Montellano, The DosS-DosT/DosR Mycobacterial Sensor System. Biosensors (Basel), 2013. 3(3): p. 259–282.

14. Xu, Y., Z. Zhao, W. Tong, Y. Ding, B. Liu, Y. Shi, J. Wang, S. Sun, M. Liu, Y. Wang, et al., An acid-tolerance response system protecting exponentially growing Escherichia coli. Nat Commun, 2020. 11(1): p. 1496.

15. Perron, K., O. Caille, C. Rossier, C. Van Delden, J.L. Dumas, and T. Kohler, CzcR-CzcS, a two-component system involved in heavy metal and carbapenem resistance in Pseudomonas aeruginosa. J Biol Chem, 2004. 279(10): p. 8761–8.

16. Yuan, J., F. Jin, T. Glatter, and V. Sourjik, Osmosensing by the bacterial PhoQ/PhoP two-component system. Proc Natl Acad Sci U S A, 2017. 114(50): p. E10792–E10798.

17. Tierney, A.R. and P.N. Rather, Roles of two-component regulatory systems in antibiotic resistance. Future Microbiol, 2019. 14: p. 533–552.

18. Jansen, A., M. Turck, C. Szekat, M. Nagel, I. Clever, and G. Bierbaum, Role of insertion elements and yycFG in the development of decreased susceptibility to vancomycin in Staphylococcus aureus. Int J Med Microbiol, 2007. 297(4): p. 205–15.

19. Guerrero, P., B. Collao, R. Alvarez, H. Salinas, E.H. Morales, I.L. Calderon, C.P. Saavedra, and F. Gil, Salmonella enterica serovar Typhimurium BaeSR two-component system positively regulates sodA in response to ciprofloxacin. Microbiology (Reading), 2013. 159(Pt 10): p. 2049–2057.

20. Bader, M.W., S. Sanowar, M.E. Daley, A.R. Schneider, U. Cho, W. Xu, R.E. Klevit, H. Le Moual, and S.I. Miller, Recognition of antimicrobial peptides by a bacterial sensor kinase. Cell, 2005. 122(3): p. 461–72.

21. Yadavalli, S.S., J.N. Carey, R.S. Leibman, A.I. Chen, A.M. Stern, M. Roggiani, A.M. Lippa, and M. Goulian, Antimicrobial peptides trigger a division block in Escherichia coli through stimulation of a signalling system. Nat Commun, 2016. 7: p. 12340.

22. Gunn, J.S., The Salmonella PmrAB regulon: lipopolysaccharide modifications, antimicrobial peptide resistance and more. Trends Microbiol, 2008. 16(6): p. 284–90.

23. Nguyen, H.T., K.A. Wolff, R.H. Cartabuke, S. Ogwang, and L. Nguyen, A lipoprotein modulates activity of the MtrAB two-component system to provide intrinsic multidrug resistance, cytokinetic control and cell wall homeostasis in Mycobacterium. Mol Microbiol, 2010. 76(2): p. 348–64.

24. Comenge, Y., R. Quintiliani, Jr., L. Li, L. Dubost, J.P. Brouard, J.E. Hugonnet, and M. Arthur, The CroRS two-component regulatory system is required for intrinsic beta-lactam resistance in Enterococcus faecalis. J Bacteriol, 2003. 185(24): p. 7184–92.

25. Bem, A.E., N. Velikova, M.T. Pellicer, P. Baarlen, A. Marina, and J.M. Wells, Bacterial histidine kinases as novel antibacterial drug targets. ACS Chem Biol, 2015. 10(1): p. 213–24.

26. Shi, X., S. Wegener-Feldbrugge, S. Huntley, N. Hamann, R. Hedderich, and L. Sogaard-Andersen, Bioinformatics and experimental analysis of proteins of two-component systems in Myxococcus xanthus. J Bacteriol, 2008. 190(2): p. 613–24.

27. Perez, J.C., D. Shin, I. Zwir, T. Latifi, T.J. Hadley, and E.A. Groisman, Evolution of a bacterial regulon controlling virulence and Mg(2+) homeostasis. PLoS Genet, 2009. 5(3): p. e1000428.

28. Palmer, A.D., K. Kim, and J.M. Slauch, PhoP-Mediated Repression of the SPI1 Type 3 Secretion System in Salmonella enterica Serovar Typhimurium. J Bacteriol, 2019. 201(16).

29. Monsieurs, P., S. De Keersmaecker, W.W. Navarre, M.W. Bader, F. De Smet, M. McClelland, F.C. Fang, B. De Moor, J. Vanderleyden, and K. Marchal, Comparison of the PhoPQ regulon in Escherichia coli and Salmonella typhimurium. J Mol Evol, 2005. 60(4): p. 462–74.

30. Schmidl, S.R., F. Ekness, K. Sofjan, K.N. Daeffler, K.R. Brink, B.P. Landry, K.P. Gerhardt, N. Dyulgyarov, R.U. Sheth, and J.J. Tabor, Rewiring bacterial two-component systems by modular DNA-binding domain swapping. Nat Chem Biol, 2019. 15(7): p. 690–698.

31. Toprak, E., A. Veres, J.B. Michel, R. Chait, D.L. Hartl, and R. Kishony, Evolutionary paths to antibiotic resistance under dynamically sustained drug selection. Nat Genet, 2011. 44(1): p. 101–5.

32. Watson, M., J.W. Liu, and D. Ollis, Directed evolution of trimethoprim resistance in Escherichia coli. FEBS J, 2007. 274(10): p. 2661–71.

33. Matange, N., S. Bodkhe, M. Patel, and P. Shah, Trade-offs with stability modulate innate and mutationally acquired drug resistance in bacterial dihydrofolate reductase enzymes. Biochem J, 2018. 475(12): p. 2107–2125.

34. Yadavalli, S.S., T. Goh, J.N. Carey, G. Malengo, S. Vellappan, B.E. Nickels, V. Sourjik, M. Goulian, and J. Yuan, Functional determinants of a small protein controlling a broadly conserved bacterial sensor kinase. J Bacteriol, 2020.

35. Lippa, A.M. and M. Goulian, Feedback inhibition in the PhoQ/PhoP signaling system by a membrane peptide. PLoS Genet, 2009. 5(12): p. e1000788.

36. Salazar, M.E., A.I. Podgornaia, and M.T. Laub, The small membrane protein MgrB regulates PhoQ bifunctionality to control PhoP target gene expression dynamics. Mol Microbiol, 2016. 102(3): p. 430–445.

37. Minagawa, S., H. Ogasawara, A. Kato, K. Yamamoto, Y. Eguchi, T. Oshima, H. Mori, A. Ishihama, and R. Utsumi, Identification and molecular characterization of the Mg2+ stimulon of Escherichia coli. J Bacteriol, 2003. 185(13): p. 3696–702.

38. Xu, J., T. Li, Y. Gao, J. Deng, and J. Gu, MgrB affects the acid stress response of Escherichia coli by modulating the expression of iraM. FEMS Microbiol Lett, 2019. 366(11).

39. Flensburg, J. and O. Skold, Massive overproduction of dihydrofolate reductase in bacteria as a response to the use of trimethoprim. Eur J Biochem, 1987. 162(3): p. 473–6.

40. Matange, N., S. Hegde, and S. Bodkhe, Adaptation Through Lifestyle Switching Sculpts the Fitness Landscape of Evolving Populations: Implications for the Selection of Drug-Resistant Bacteria at Low Drug Pressures. Genetics, 2019. 211(3): p. 1029–1044.

41. Wistrand-Yuen, E., M. Knopp, K. Hjort, S. Koskiniemi, O.G. Berg, and D.I. Andersson, Evolution of high-level resistance during low-level antibiotic exposure. Nat Commun, 2018. 9(1): p. 1599.

42. Zhou, J., Y. Dong, X. Zhao, S. Lee, A. Amin, S. Ramaswamy, J. Domagala, J.M. Musser, and K. Drlica, Selection of antibiotic-resistant bacterial mutants: allelic diversity among fluoroquinolone-resistant mutations. J Infect Dis, 2000. 182(2): p. 517–25.

43. Nagakubo, S., K. Nishino, T. Hirata, and A. Yamaguchi, The putative response regulator BaeR stimulates multidrug resistance of Escherichia coli via a novel multidrug exporter system, MdtABC. J Bacteriol, 2002. 184(15): p. 4161–7.

44. Eguchi, Y., E. Ishii, K. Hata, and R. Utsumi, Regulation of acid resistance by connectors of two-component signal transduction systems in Escherichia coli. J Bacteriol, 2011. 193(5): p. 1222–8.

45. Eguchi, Y., J. Itou, M. Yamane, R. Demizu, F. Yamato, A. Okada, H. Mori, A. Kato, and R. Utsumi, B1500, a small membrane protein, connects the two-component systems EvgS/EvgA and PhoQ/PhoP in Escherichia coli. Proc Natl Acad Sci U S A, 2007. 104(47): p. 18712–7.

46. Iwase, T., T. Matsuo, S. Nishioka, A. Tajima, and Y. Mizunoe, Hydrophobicity of Residue 128 of the Stress-Inducible Sigma Factor RpoS Is Critical for Its Activity. Front Microbiol, 2017. 8: p. 656.

47. Smith, D.R., J.E. Price, P.E. Burby, L.P. Blanco, J. Chamberlain, and M.R. Chapman, The Production of Curli Amyloid Fibers Is Deeply Integrated into the Biology of Escherichia coli. Biomolecules, 2017. 7(4).

48. Andersson, D.I. and D. Hughes, Microbiological effects of sublethal levels of antibiotics. Nat Rev Microbiol, 2014. 12(7): p. 465–78.

49. Gullberg, E., S. Cao, O.G. Berg, C. Ilback, L. Sandegren, D. Hughes, and D.I. Andersson, Selection of resistant bacteria at very low antibiotic concentrations. PLoS Pathog, 2011. 7(7): p. e1002158.

50. Matange, N., Highly Contingent Phenotypes of Lon Protease Deficiency in Escherichia coli upon Antibiotic Challenge. J Bacteriol, 2020. 202(3).

51. Groisman, E.A., The pleiotropic two-component regulatory system PhoP-PhoQ. J Bacteriol, 2001. 183(6): p. 1835–42.

52. Lippa, A.M. and M. Goulian, Perturbation of the oxidizing environment of the periplasm stimulates the PhoQ/PhoP system in Escherichia coli. J Bacteriol, 2012. 194(6): p. 1457–63.

53. Mmatli, M., N.M. Mbelle, N.E. Maningi, and J. Osei Sekyere, Emerging Transcriptional and Genomic Mechanisms Mediating Carbapenem and Polymyxin Resistance in Enterobacteriaceae: a Systematic Review of Current Reports. mSystems, 2020. 5(6).

54. Cannatelli, A., M.M. D’Andrea, T. Giani, V. Di Pilato, F. Arena, S. Ambretti, P. Gaibani, and G.M. Rossolini, In vivo emergence of colistin resistance in Klebsiella pneumoniae producing KPC-type carbapenemases mediated by insertional inactivation of the PhoQ/PhoP mgrB regulator. Antimicrob Agents Chemother, 2013. 57(11): p. 5521–6.

55. Haeili, M., A. Javani, J. Moradi, Z. Jafari, M.M. Feizabadi, and E. Babaei, MgrB Alterations Mediate Colistin Resistance in Klebsiella pneumoniae Isolates from Iran. Front Microbiol, 2017. 8: p. 2470.

56. Silva, D.M.D., C. Faria-Junior, D.R. Nery, P.M. Oliveira, L.O.R. Silva, E.G. Alves, G. Lima, and A.L. Pereira, Insertion sequences disrupting mgrB in carbapenem-resistant Klebsiella pneumoniae strains in Brazil. J Glob Antimicrob Resist, 2021. 24: p. 53–57.

57. Kidd, T.J., G. Mills, J. Sa-Pessoa, A. Dumigan, C.G. Frank, J.L. Insua, R. Ingram, L. Hobley, and J.A. Bengoechea, A Klebsiella pneumoniae antibiotic resistance mechanism that subdues host defences and promotes virulence. EMBO Mol Med, 2017. 9(4): p. 430–447.

58. Baym, M., T.D. Lieberman, E.D. Kelsic, R. Chait, R. Gross, I. Yelin, and R. Kishony, Spatiotemporal microbial evolution on antibiotic landscapes. Science, 2016. 353(6304): p. 1147–51.

59. Levin-Reisman, I., I. Ronin, O. Gefen, I. Braniss, N. Shoresh, and N.Q. Balaban, Antibiotic tolerance facilitates the evolution of resistance. Science, 2017. 355(6327): p. 826–830.

60. Yang, J., Y. Ogawa, H. Camakaris, T. Shimada, A. Ishihama, and A.J. Pittard, folA, a new member of the TyrR regulon in Escherichia coli K-12. J Bacteriol, 2007. 189(16): p. 6080–4.

61. Razzell, W.E. and P. Casshyap, Substrate Specificity and Induction of Thymidine Phosphorylase in Escherichia Coli. J Biol Chem, 1964. 239: p. 1789–93.

62. Rodrigues, J.V. and E.I. Shakhnovich, Adaptation to mutational inactivation of an essential gene converges to an accessible suboptimal fitness peak. Elife, 2019. 8.

63. Erzberger, J.P., M.L. Mott, and J.M. Berger, Structural basis for ATP-dependent DnaA assembly and replication-origin remodeling. Nat Struct Mol Biol, 2006. 13(8): p. 676–83.

64. Kawakami, H., S. Ozaki, S. Suzuki, K. Nakamura, T. Senriuchi, M. Su’etsugu, K. Fujimitsu, and T. Katayama, The exceptionally tight affinity of DnaA for ATP/ADP requires a unique aspartic acid residue in the AAA+ sensor 1 motif. Mol Microbiol, 2006. 62(5): p. 1310–24.

65. Guzman, E.C. and C.M. Martin, Thymineless death, at the origin. Front Microbiol, 2015. 6: p. 499.

66. Hicks, N.D., S.R. Giffen, P.H. Culviner, M.C. Chao, C.L. Dulberger, Q. Liu, S. Stanley, J. Brown, J. Sixsmith, I.D. Wolf, et al., Mutations in dnaA and a cryptic interaction site increase drug resistance in Mycobacterium tuberculosis. PLoS Pathog, 2020. 16(11): p. e1009063.

67. Tsilibaris, V., G. Maenhaut-Michel, and L. Van Melderen, Biological roles of the Lon ATP-dependent protease. Res Microbiol, 2006. 157(8): p. 701–13.

68. Cole, S.T., Characterisation of the promoter for the LexA regulated sulA gene of Escherichia coli. Mol Gen Genet, 1983. 189(3): p. 400–4.

69. Gottesman, S., E. Halpern, and P. Trisler, Role of sulA and sulB in filamentation by lon mutants of Escherichia coli K-12. J Bacteriol, 1981. 148(1): p. 265–73.

70. Spitz, F. and D. Duboule, Global control regions and regulatory landscapes in vertebrate development and evolution. Adv Genet, 2008. 61: p. 175–205.

71. Bayer, T.S., Using synthetic biology to understand the evolution of gene expression. Curr Biol, 2010. 20(17): p. R772–9.

72. Rodrigues, J.V., S. Bershtein, A. Li, E.R. Lozovsky, D.L. Hartl, and E.I. Shakhnovich, Biophysical principles predict fitness landscapes of drug resistance. Proc Natl Acad Sci U S A, 2016. 113(11): p. E1470–8.

73. Gilad, Y., A. Oshlack, and S.A. Rifkin, Natural selection on gene expression. Trends Genet, 2006. 22(8): p. 456–61.

74. Signor, S.A. and S.V. Nuzhdin, The Evolution of Gene Expression in cis and trans. Trends Genet, 2018. 34(7): p. 532–544.

75. Thompson, D., A. Regev, and S. Roy, Comparative analysis of gene regulatory networks: from network reconstruction to evolution. Annu Rev Cell Dev Biol, 2015. 31: p. 399–428.

76. Dong, T., S.M. Chiang, C. Joyce, R. Yu, and H.E. Schellhorn, Polymorphism and selection of rpoS in pathogenic Escherichia coli. BMC Microbiol, 2009. 9: p. 118.

77. Kotewicz, M.L., E.W. Brown, J. Eugene LeClerc, and T.A. Cebula, Genomic variability among enteric pathogens: the case of the mutS-rpoS intergenic region. Trends Microbiol, 2003. 11(1): p. 2–6.

78. Martinez-Garcia, E., A. Tormo, and J.M. Navarro-Llorens, Polymorphism in the yclC-rpoS region of enterobacteria. Curr Microbiol, 2003. 46(5): p. 365–70.

79. Wrande, M., J.R. Roth, and D. Hughes, Accumulation of mutants in “aging” bacterial colonies is due to growth under selection, not stress-induced mutagenesis. Proc Natl Acad Sci U S A, 2008. 105(33): p. 11863–8.

80. Chiang, S.M., T. Dong, T.A. Edge, and H.E. Schellhorn, Phenotypic diversity caused by differential RpoS activity among environmental Escherichia coli isolates. Appl Environ Microbiol, 2011. 77(22): p. 7915–23.

81. Liu, B., G. Eydallin, R.P. Maharjan, L. Feng, L. Wang, and T. Ferenci, Natural Escherichia coli isolates rapidly acquire genetic changes upon laboratory domestication. Microbiology (Reading), 2017. 163(1): p. 22–30.

82. Baquero, F., Low-level antibacterial resistance: a gateway to clinical resistance. Drug Resist Updat, 2001. 4(2): p. 93–105.

83. Gutierrez, A., L. Laureti, S. Crussard, H. Abida, A. Rodriguez-Rojas, J. Blazquez, Z. Baharoglu, D. Mazel, F. Darfeuille, J. Vogel, et al., beta-Lactam antibiotics promote bacterial mutagenesis via an RpoS-mediated reduction in replication fidelity. Nat Commun, 2013. 4: p. 1610.

84. Hiltunen, T., M. Virta, and A.L. Laine, Antibiotic resistance in the wild: an eco-evolutionary perspective. Philos Trans R Soc Lond B Biol Sci, 2017. 372(1712).

85. Bjorkman, J., I. Nagaev, O.G. Berg, D. Hughes, and D.I. Andersson, Effects of environment on compensatory mutations to ameliorate costs of antibiotic resistance. Science, 2000. 287(5457): p. 1479–82.

86. Handel, A., R.R. Regoes, and R. Antia, The role of compensatory mutations in the emergence of drug resistance. PLoS Comput Biol, 2006. 2(10): p. e137.

87. Melnyk, A.H., A. Wong, and R. Kassen, The fitness costs of antibiotic resistance mutations. Evol Appl, 2015. 8(3): p. 273–83.

88. Qi, Q., M. Toll-Riera, K. Heilbron, G.M. Preston, and R.C. MacLean, The genomic basis of adaptation to the fitness cost of rifampicin resistance in Pseudomonas aeruginosa. Proc Biol Sci, 2016. 283(1822).

89. Schulz zur Wiesch, P., J. Engelstadter, and S. Bonhoeffer, Compensation of fitness costs and reversibility of antibiotic resistance mutations. Antimicrob Agents Chemother, 2010. 54(5): p. 2085–95.

90. Baba, T., T. Ara, M. Hasegawa, Y. Takai, Y. Okumura, M. Baba, K.A. Datsenko, M. Tomita, B.L. Wanner, and H. Mori, Construction of Escherichia coli K-12 in-frame, single-gene knockout mutants: the Keio collection. Mol Syst Biol, 2006. 2: p. 2006 0008.

91. Yamamoto, N., K. Nakahigashi, T. Nakamichi, M. Yoshino, Y. Takai, Y. Touda, A. Furubayashi, S. Kinjyo, H. Dose, M. Hasegawa, et al., Update on the Keio collection of Escherichia coli single-gene deletion mutants. Mol Syst Biol, 2009. 5: p. 335.

92. Datsenko, K.A. and B.L. Wanner, One-step inactivation of chromosomal genes in Escherichia coli K-12 using PCR products. Proc Natl Acad Sci U S A, 2000. 97(12): p. 6640–5.

93. Deatherage, D.E. and J.E. Barrick, Identification of mutations in laboratory-evolved microbes from next-generation sequencing data using breseq. Methods Mol Biol, 2014. 1151: p. 165–88.

94. Kitagawa, M., T. Ara, M. Arifuzzaman, T. Ioka-Nakamichi, E. Inamoto, H. Toyonaga, and H. Mori, Complete set of ORF clones of Escherichia coli ASKA library (a complete set of E. coli K-12 ORF archive): unique resources for biological research. DNA Res, 2005. 12(5): p. 291–9.

