## Supplementary figures for "Adaptation and Compensation in a Bacterial Gene Regulatory Network Evolving Under Antibiotic Selection"

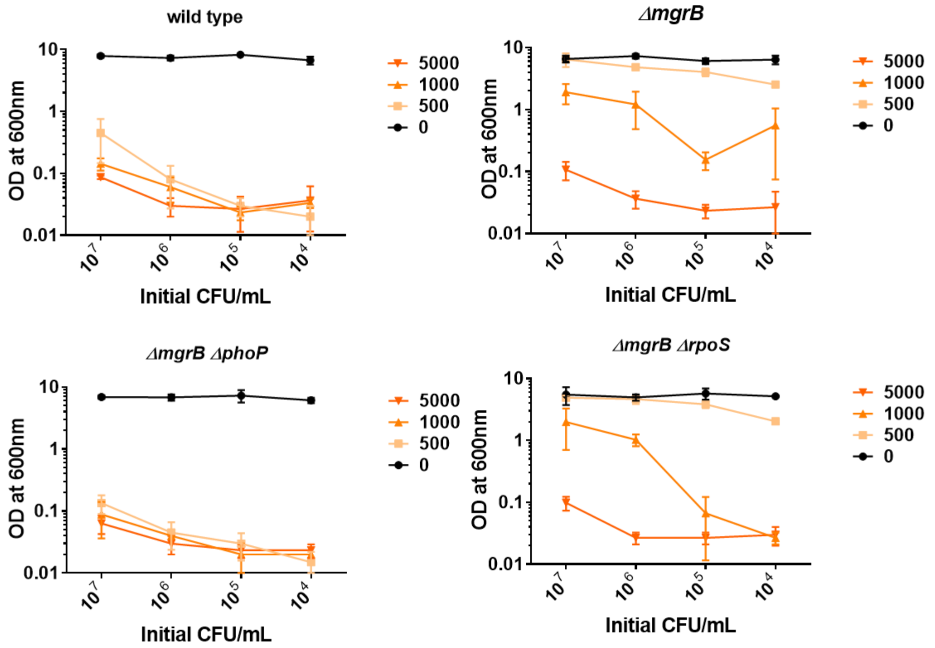


**Figure S1.** Colonization of indicated *E. coli* strains in growth media supplemented with increasing trimethoprim concentrations (5000, 1000, 500, 0 ng/mL) starting at the indicated cell densities (X-axis). Optical density of cultures was measured after 24 hours of growth. Mean ± S.D. are plotted.

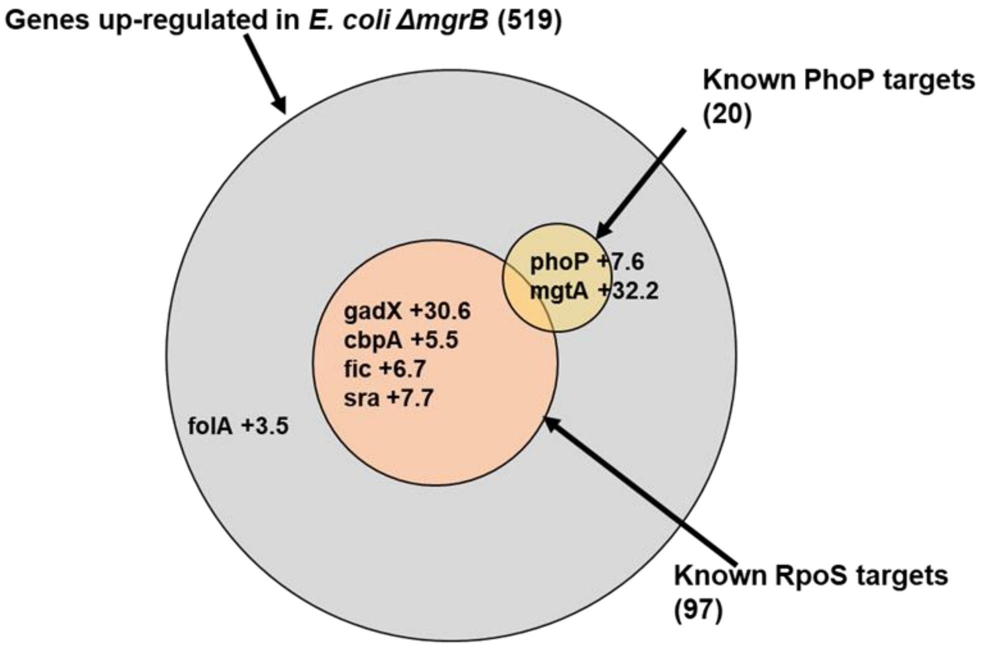


**Figure S2.** Diagrammatic representation of up-regulated genes in *E. coli ΔmgrB*. Transcriptomic data for *E. coli ΔmgrB* was taken from Xu et al. (2019) [1]. Area of each of the circles in the Venn diagram is proportional to the number of genes in that set. PhoP and RpoS regulated genes were determined using RegulonDB [2]. Representative genes or those relevant to this study and their level of up-regulation are shown.

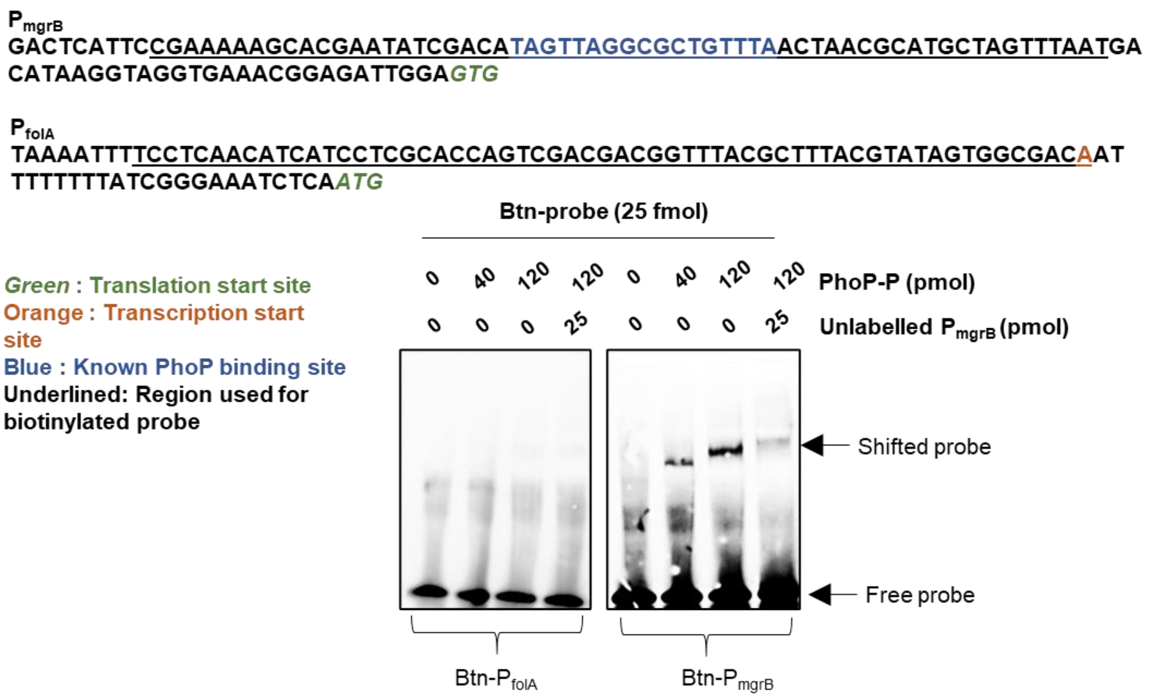


**Figure S3.** Electrophoretic mobility shift assay (EMSA) to test *in vitro* binding of phosphorylated PhoP (PhoP-P) to the promoter of *folA* ($P_{folA}$). Promoter of *mgrB* ($P_{mgrB}$) was used as positive control [3]. Sequences of the 2 promoters are shown. The probe consisted of 60-mer oligonucleotides (underlined) with biotinylation at the 5' end. The amounts of purified PhoP-P, labelled and unlabelled competitor used for each reaction are indicated. Positions of free and shifted probes are indicated. No shift was detectable for $P_{folA}$.

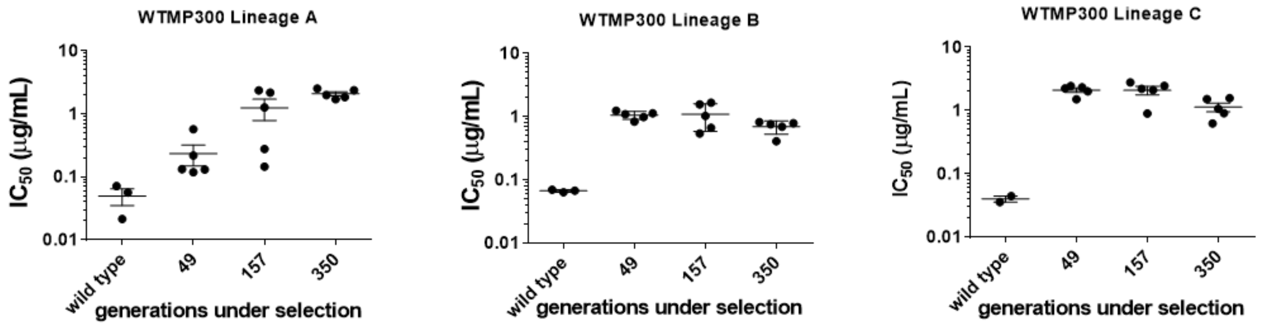


**Figure S4.** IC50 values of trimethoprim for resistant isolates from WTMP300 lineages at 47, 157 and 350 generations. Each spot represents mean IC50 from 2 independent measurements for an individual isolate. Mean and S.E.M. of 4-6 isolates from each time point is shown. Ancestral IC50 (wild type) from three replicate measurements and mean ± S.E.M. are shown.

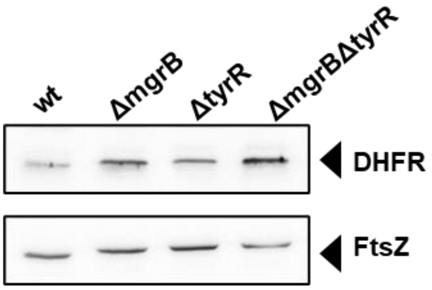

**Figure S5.** DHFR protein levels measured in wild type and mutant strains by immunoblotting with anti-DHFR antibody. Anti-FtsZ was used as a loading control.

***E. coli K-12 MG1655***
VKKFRWVVLVVVVLACLLLWAQVFNMMCDQDVQFFSGICAINQFIPW

**TMPR5, *E. coli* strain NCTC9075**
VKKFRWVVLVVVVLACLLLWAQVFNMMCDQDVQFSAEFVPLTSLSRGDIIFSALDFLPLQQW

**WTMP50 isolates**
VKKFRWVVLVVVVLACLLLWAQVFNMMCDQDVHFSAEFVPLTSLSRGDIIFSALDFLPLQQW

**LTMPR300 isolates**
VKKFRWVVLVVVVLACLLLWAQVFNMMCDTGCTIFQRNLCH

***E. coli* CVM N18EC0244, CVM N17EC0040, CVM N17EC0046, NCTC9029**
VKKFRWVALVVVVLACLLLWAQVFNMMCDQDVQFFSGICAINQFIRGDIIFSALDFLPLQQW

***E. coli* CVM N20EC1291**
VKKFRWVALVVVVLACLLLWAQVFNMMCDQDVQFFSGICAINQFIPWCIIFSALDFLPLQQW

***E. coli* SJP130** VKKFRWVALVVVVLACLLLWAQVFNMMCDQDVQFFSGICAINQFIPGDIIFSALDFLPLQQW

***E. coli* VREC0557**
VKKFRWVALVVVVLACLLLWAQVFNMMCDQDVQFFSGICAINQFIPWWVMTPTY

***E. coli* ST73**
VKKFRWVALVVVVLACLLLWAQVFNMMCDQDVQFFSGICAINQFIPWRYHFFSTRFSSSPAVVE

Green : cytosolic
Orange : trans-membrane
Blue : periplasmic (underlined residues are required for activity)
Black : altered sequenced in mutants

| *E. coli* Strain | Accession number | Source / Collection / Other available information | Mutation in mgrB | Reference |
|---|---|---|---|---|
| NCTC 9075 | UGEM0100000 4.1 | WTSI, Pathogen Informatics (U.K.) Serotype 075:K:H5 | ΔT (+100) | - |
| CVM N18EC0244 | AARKPQ0100 00008.1 | NARMS (CDC, U.S.A) Isolated from retail meat | ΔC (+135) | Tyson et al. (2019) |
| CVM N17EC0040 | RNLX0100001 4.1 | NARMS (CDC, U.S.A) Isolated from retail meat | ΔC (+135) | Tyson et al. (2019) |
| CVM N17EC0046 | RNLY0100000 6.1 | NARMS (CDC, U.S.A) Isolated from retail meat | ΔC (+135) | Tyson et al. (2019) |
| NCTC 9029 | UGDE0100000 1.1 | WTSI, Pathogen Informatics (U.K.) Serotype 029:H10 | ΔC (+135) | - |
| CVM N20EC1291 | AAZRQZ01000 0006.1 | NARMS (CDC, U.S.A) Isolated from retail meat | A→T (+144) | Tyson et al. (2019) |
| SJP130 | DADPHN0100 00001.1 | NCBI Pathogen Detection Project | ΔG (+138) | Souvarov et al. (2018) |
| VREC0557 | NZ_UINQ0100 0006.1 | WTSI, Pathogen Informatics (U.K.) Isolated from *Meleagris gallopavo* faeces | A→G (+144) | - |
| ST73 | NZ_UNQU010 00003.1 | Clinical ExPEC sourced from Sydney Isolated from Mid-stream urine of diseased human | T→C (+142) | - |

**Figure S6.** Predicted protein sequences of mutant MgrB alleles identified in this study, and from publicly available sequences of pathogenic and environmental *E. coli* strains identified by BLAST analysis. Each sequence is coloured based on the functional annotation of MgrB. Accession numbers and source of strain [4, 5] are indicated in the table.

*Salmonella enterica* subsp. *enterica* serovar Typhimurium isolate AUSMDU00003672
VKKFRWVVLGIVVVVCLLLWAQVFNIMCDQDVQFFSGICAINKFIPWYTFYTRDFLPV

*Salmonella enterica* subsp. *enterica* serovar Typhi strain 1071_07
VKKFRWVVLGIVVVVCLLLWAQVFNIMCDQDVQFFQRYLRHSINFIPW

*Salmonella enterica* subsp. *enterica* NCTC 6245
VKKFRWVVLGIVVVVCLLLWAQVFNIMCDQDVQFFSGICAINKFIPGKRFTPEISFPCSSGRMTSLH

*Citrobacter freundii 2021DK-00032*
MKKIRWVVLIVVVLVCILMWAQVFNIMCDQDVQFFSGICAINKFIPWLAHFCHS

Green : cytosolic
Orange : trans-membrane
Blue : periplasmic
Black : predicted altered sequenced in mutants

| Strain | Accession number | Source / Collection / Other available information | Reference |
|---|---|---|---|
| *Salmonella enterica* | | | |
| serovar Typhimurium isolate AUSMDU00003672 | AAMVMG010000117 | GenomeTrakr network<br>Isolate from food-borne Salmonella outbreak | - |
| serovar Typhi strain 1071_07 | QAVT01000062.1 | Isolate from an outbreak of Salmonella (Brazil) | - |
| NCTC 6245 | UGXL01000002.1 | WTSI, Pathogen Informatics (U.K.)<br>serotype Manhattan | - |
| *Citrobacter freundii* | | | |
| 2021DK-00032 | ABAGBY010000005 | Clinical and Environmental Microbiology Branch: Whole genome sequencing antimicrobial resistance pathogens in the healthcare setting<br><br>Isolated from a rectal swab from human | - |

**Figure S7.** Predicted protein sequences of MgrB alleles from publicly available sequences of pathogenic and environmental *S. typhimurium* and *C. freundii* strains identified by BLAST analysis. Each sequence is coloured based on the functional annotation of MgrB for *E. coli*. Accession numbers and source of strain are indicated in the table.

**Table S1. Summary of genetic changes in trimethoprim resistant isolates from WTMP300 Lineage A identified by genome sequencing**

**Table S2. Summary of genetic changes in trimethoprim resistant and tolerant isolates from WTMP50 Lineage A by genome sequencing**

**Table S3. Summary of genetic changes in trimethoprim resistant isolates from LTMP300 Lineage A by genome sequencing**

**(Table S1, S2 and S3 are separate files)**

**Supplementary references**


1. Xu, J., T. Li, Y. Gao, J. Deng, and J. Gu, *MgrB affects the acid stress response of Escherichia coli by modulating the expression of iraM.* FEMS Microbiol Lett, 2019. **366**(11).
2. Gama-Castro, S., H. Salgado, A. Santos-Zavaleta, D. Ledezma-Tejeida, L. Muniz-Rascado, J.S. Garcia-Sotelo, K. Alquicira-Hernandez, I. Martinez-Flores, L. Pannier, J.A. Castro-Mondragon, et al., *RegulonDB version 9.0: high-level integration of gene regulation, coexpression, motif clustering and beyond.* Nucleic Acids Res, 2016. **44**(D1): p. D133-43.
3. Yamamoto, K., H. Ogasawara, N. Fujita, R. Utsumi, and A. Ishihama, *Novel mode of transcription regulation of divergently overlapping promoters by PhoP, the regulator of two-component system sensing external magnesium availability.* Mol Microbiol, 2002. **45**(2): p. 423-38.
4. Souvorov, A., R. Agarwala, and D.J. Lipman, *SKESA: strategic k-mer extension for scrupulous assemblies.* Genome Biol, 2018. **19**(1): p. 153.
5. Tyson, G.H., C. Li, C.H. Hsu, S. Bodeis-Jones, and P.F. McDermott, *Diverse Fluoroquinolone Resistance Plasmids From Retail Meat E. coli in the United States.* Front Microbiol, 2019. **10**: p. 2826.
